# Depletion and replacement of tissue resident macrophages in mice with germ-line deletion of a conserved enhancer in the *Csf1r* locus

**DOI:** 10.64898/2026.03.22.713539

**Authors:** Yajun Liu, Sebastien Jacquelin, Isis Taylor, Emma K. Green, Omkar L. Patkar, Sahar Keshvari, Ginell Ranpura, Conan J. O. O’Brien, Eline Jessen, Emma Maxwell, Rachel Allavena, Alexandre Gallerand, Stoyan Ivanov, Antony Adamson, Neil E. Humphreys, Kim M. Summers, Katharine M. Irvine, David A. Hume

**Affiliations:** Mater Research Institute-University of Queensland, Translational Research Institute, Woolloongabba, Brisbane, Qld 4102, Australia; School of Veterinary Science, The University of Queensland, Gatton, Qld 4343, Australia; Université Côte d’Azur, CNRS, LP2M, Nice, France; Faculty of Biology, Medicine and Health, University of Manchester, Manchester, UK

**Keywords:** CSF1R, monocyte, macrophage, enhancer, transgenic mouse, quantitative signalling

## Abstract

Expression of the *Csf1r* gene in cells of the mononuclear phagocyte lineage is regulated by a conserved enhancer, the fms-intronic regulatory element (FIRE). In mice with a germ-line deletion of FIRE (*Fireko*) CSF1R expression is undetectable in bone marrow progenitors and classical monocytes. *Fireko* mice lack subpopulations of macrophages in the brain and periphery but develop normally. Here we show that loss of CSF1R expression in *Fireko* mice is partly overcome by CSF2 *in vitro* and inflammatory recruitment *in vitro.* Analysis of heterozygous mutant mice and deletion of the conserved AP1 motif in FIRE provide evidence that continuous receptor synthesis determines CSF1 responsiveness. The absence of macrophages in kidney and heart of *Fireko* mice was not associated with detectable loss of physiological function. In a model of renal injury macrophage recruitment and histopathology were similar in WT and *Fireko* mice. Tissue resident macrophages that were depleted in *Fireko* mice, including microglia, were replaced by donor-derived cells following intraperitoneal adoptive transfer of wild-type bone marrow at weaning. The *Fireko* mouse provides a novel platform to dissect the functions of tissue resident macrophages in development, homeostasis and pathology.

**Summary Statement:** This study describes a unique model of selective tissue resident macrophage deficiency arising from dysregulated expression of the mouse *Csf1r* gene.

## Introduction

The mononuclear phagocyte system (MPS) is a family of cells including progenitors in bone marrow, circulating blood monocytes and resident tissue macrophages in every organ in the body (Grabert et al., 2020; Hume et al., 2019). Proliferation, differentiation and survival of cells of the MPS depends upon signals from the macrophage colony-stimulating factor receptor, CSF1R (Chitu and Stanley, 2017; Hume et al., 2020). *Csf1r^-/-^* mice (Dai et al., 2002), rats (Carter-Cusack et al., 2024; Keshvari et al., 2021; Pridans et al., 2018) and humans (Guo et al., 2019; Guo and Ikegawa, 2021; Oosterhof et al., 2019) lack bone-resorbing osteoclasts in bone and have reduced density of resident tissue macrophages in most organs. In the rodent models, *Csf1r^-/-^* pups are indistinguishable from litter mates at birth but postnatal skeletal development, somatic growth, organ maturation and fertility are severely compromised leading to early mortality (Carter-Cusack et al., 2024; Chitu and Stanley, 2017; Keshvari et al., 2021). On a pure C57BL/6J mouse background, homozygous *Csf1r* null mutation leads to perinatal lethality (Percin et al., 2018) whilst on the FVB/J background few pups survive to weaning due to hydrocephalus (Erblich et al., 2011).

We recently described a hypomorphic mutation in the mouse *Csf1r* gene (Rojo et al., 2019). These mice harbor a deletion of a 300bp enhancer region (FIRE) that is conserved from reptiles and birds to humans (Hume et al., 2017). Homozygous mutant mice lack macrophages in the embryonic yolk sac and fetal liver (Munro et al., 2020) but unlike *Csf1r*^-/-^ mice (Dai et al., 2002), *Csf1r*^ΔFIRE/ΔFIRE^ (hereafter *Fireko*) mice were not osteoclast-deficient/osteopetrotic or growth-inhibited and were viable and fertile as adults (Rojo et al., 2019). The analysis of *Fireko* mice has focussed mainly on the impacts of microglial deficiency (reviewed in (Hume, 2025)). *Fireko* mice also lack resident macrophages in the peritoneal cavity and omentum, epidermis, kidney, heart and salivary gland (Louwe et al., 2022; McKendrick et al., 2023; Rojo et al., 2019), whereas in other organs (liver, spleen, intestine, lung) CSF1R-dependent resident macrophages appeared unaffected and *Csf1r* mRNA was expressed normally. The molecular basis for the selective impact of the *Fireko* is unknown.

Like homozygous *Csf1r* or *Csf1* mutation (Chitu and Stanley, 2017; Percin et al., 2018), the *Fireko* was lethal on the C57BL/6J background (Preprint; (Taylor et al., 2025)). The *Csf1r*^ΔFIRE^ allele was intercrossed with a *Csf1r-*EGFP transgenic line (Sasmono et al., 2003) also backcrossed to C57BL/6J. On this congenic background *Fireko* mice were identified at 59% of expected Mendelian frequency at weaning. Survival was attributed to retention of non-C57BL/6J genomic sequences despite the backcross (Preprint; (Taylor et al., 2025)). Of the homozygous pups identified around a third developed severe hydrocephalus (HC). The remainder were long-lived, healthy and fertile.

In the present study we take advantage of the congenic line to extend the investigation of the *Fireko* mouse to a more comprehensive range of peripheral tissues. We explore the mechanisms that underlie selective loss of macrophages in some tissues but not in others and the impact of heterozygous mutation on CSF1 responsiveness. We demonstrate that IP injection of wild-type marrow in *Fireko* mice at weaning leads to selective population of the vacant macrophage territories with donor-derived cells whilst making no contribution to blood monocytes.

## Results

### The loss of CSF1R in progenitors does not alter myeloid lineage commitment or monocytopoiesis

*Fireko* mice are not osteopetrotic and the mutation has no significant effect on total white cell, red cell or platelet count expressed/ml blood or on relative neutrophil, lymphocyte or monocyte counts defined by automatic analyser (Rojo et al., 2019). We first revisited expression of CSF1R and surface markers of progenitor populations and blood leukocytes on the inbred C57BL/6J congenic background. During lineage commitment, CSF1R is first expressed on the MPP3 subset of progenitors (Maxwell et al., 2026). We found no significant differences between WT and *Fireko* in the relative abundance of hematopoietic stem cells (HSC) or multipotent progenitor (MPP) subsets defined by surface markers including the growth factor receptors, KIT and FLT3 (**Figure 1A-D**, Gating strategy in **Figure S1A**). As previously reported on the original background (Rojo et al., 2019) CD115 and *Csf1r* mRNA were almost undetectable in *Fireko* marrow monocytes (**Figure 1E-F**). Mouse blood monocytes have been divided into classical and non-classical subpopulations based upon cell surface markers including Ly6C (Geissmann et al., 2010). The percentage of monocytes in the bone marrow did not distinguish *Fireko* from WT but there was a significant increase in the relative proportion of Ly6C^low^ cells (**Figure S1B**). Mouse spleen can also provide a reservoir of blood monocytes mobilized in response to inflammation (Swirski et al., 2009). As in the bone marrow, the *Fireko* did not alter the % of monocytes in spleen compared to WT but increased the proportion of Ly6C^low^ monocytes (**Figure S1C**).

**Figure 1.**
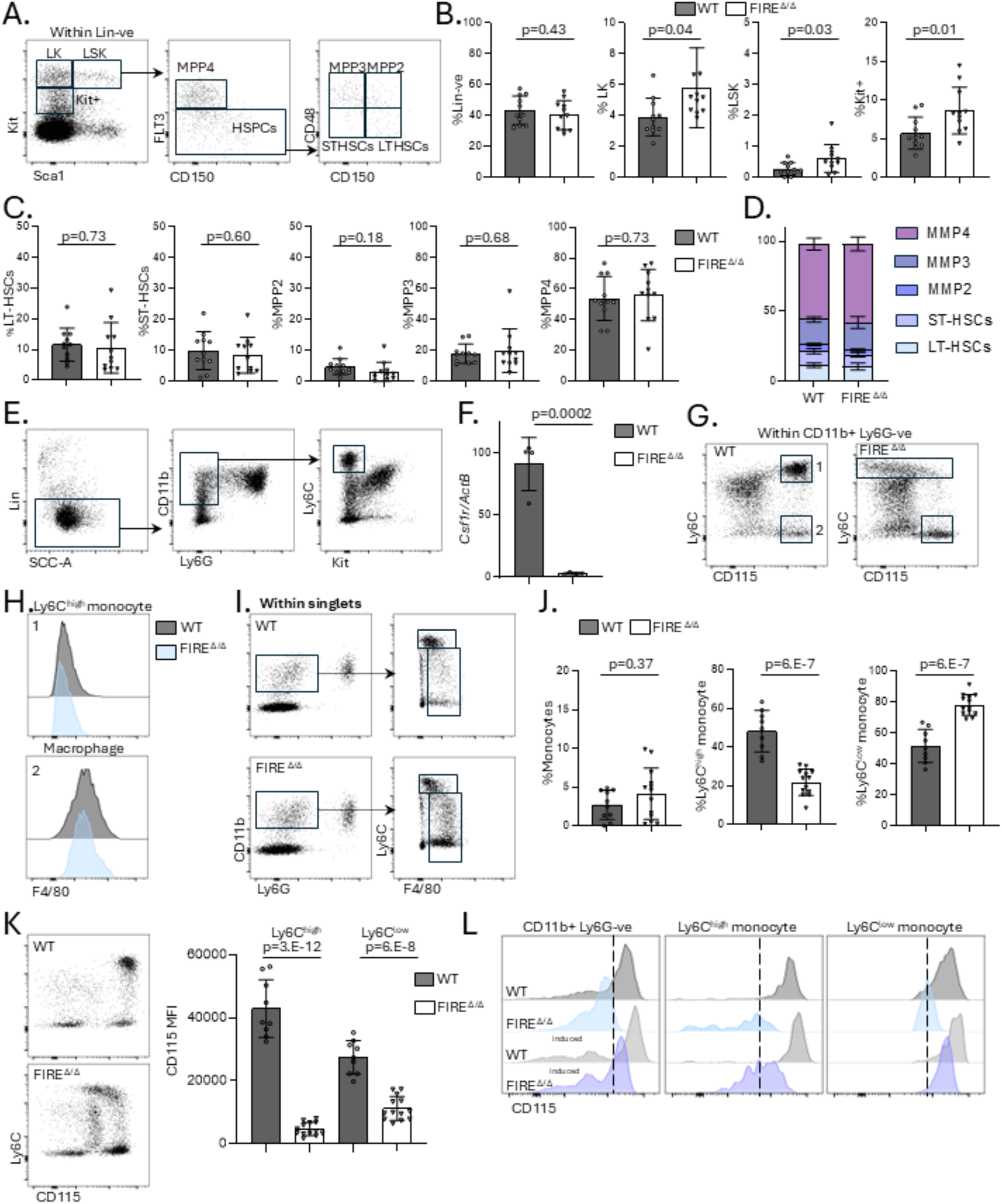
CSF1R expression and monocytopoiesis in *Fireko* mice. Bone marrow (BM) and blood cells from WT and *Fireko* (FIRE ^Δ/Δ^) mice on the inbred C57BL/6J background were isolated and analysed by flow cytometry as described in Materials and Methods. (A) BM progenitors were defined as lineage-negative (Lin-) and further divided into multipotent progenitors (MPP), long-term and short-term hematopoietic stem cells (LT-HSC, ST-HSC) based upon detection of KIT, SCA1, FLT3, CD150 and CD48 as shown in the gating strategy in Figure S1. (B) Proportions of Lin-cells (% total BM cells) and of subpopulations (% of lin-cells) in WT and FIRE ^Δ/Δ^ BM. (C, D) Proportions of LT-HSC, ST-HSC, MPP2, MPP3 and MPP4 expressed as % of LSK population. (E) BM Ly6C^high^ monocytes were defined as CD11b^+^, Ly6C^+^, Ly6G^-^, Kit^-^ as indicated in the gating strategy shown. (F) Ly6C^high^ monocytes were isolated from WT and FIRE ^Δ/Δ^ BM as shown and *Csf1r* mRNA was determined by qRT-PCR (n=4). (G) Representative flow cytometry profiles of CD115 detection in CD11b^+^, Ly6G^-^ BM cells from WT and FIRE ^Δ/Δ^ mice. (H) Representative profiles of expression of F4/80 on Ly6C^high^ monocytes and Ly6C^-^ macrophages defined in Panel G. (I) Peripheral blood monocytes were defined as Ly6G^-^, CD11b^+^, F4/80^+^ as indicated in the gating strategy shown. (J) Left panel shows quantitation of monocytes (combined boxes in Panel I) as a % of total cells in WT and FIRE ^Δ/Δ^. Right panels show proportions of Ly6C^high^ and Ly6C^low^ monocytes within the combined monocyte gate (Panel I). Each point is an individual animal. (K) Representative flow cytometry profiles and quantitation of mean fluorescence intensity (MFI) of CD115 detection in CD11b^+^, Ly6G^-^ monocytes from WT and FIRE ^Δ/Δ^ mice. (L) Figure shows representative profiles of CD115 detection in monocyte fractions defined as in Panel I. Cells were analysed with (lower panels) or without (upper panels) preincubation for 90 minutes at 37°C. Data show means and standard deviation. *p* values were determined by unpaired two-tailed t-test.

F4/80^high^ bone marrow resident macrophages associated with hematopoietic/erythroblastic islands are fragmented during tissue disaggregation and mainly appear as remnants associated with unrelated cells on flow cytometry (Millard et al., 2021). Nevertheless, CD11b^+^/Ly6C^-^/CD115^+^ cells were retained in mutant marrow (population 2 in **Figure 1G**). Unlike Ly6C^high^ bone marrow monocytes (population 1 in **Figure 1G**), these cells expressed high levels of F4/80 in both WT and *Fireko* mice consistent with their identity as resident marrow macrophages (Hume et al., 1983) (**Figure 1H**). As in marrow and spleen, the absolute monocyte count in peripheral blood was unchanged but the relative proportion of Ly6C^low^ monocytes was significantly increased (**Figure 1I-J**). shows the profile of CD115 detection in from WT and *Fireko* mice. In freshly-isolated WT monocytes the mean fluorescence intensity (MFI) of CD115 was lower in Ly6C^low^ monocytes compared to Ly6C^high^ **(Figure 1K**). In the *Fireko*, CD115 was barely detectable in the Ly6C^high^ monocytes but Ly6C^low^ monocytes retained significant expression. We reasoned that surface receptor might be masked by endogenous CSF1 in the *Fireko.* To test this possibility, leukocytes were washed and incubated at 4°C for 90 minutes *in vitro* to allow ligand dissociation prior to staining. As shown in **Figure 1L**, preincubation revealed detectable surface CD115 in *Fireko* blood monocytes, predominantly in the Ly6C^low^ cells, albeit still 5-10 fold lower than WT.

### Fireko mice are unresponsive to CSF1

To confirm the CSF1 unresponsiveness of *Fireko* mice, we injected a CSF1-Fc fusion protein, which causes expansion of monocyte numbers in marrow, monocytosis, increased tissue macrophages and growth of the liver and spleen in WT mice (Gow et al., 2014; Keshvari et al., 2024). Cohorts of WT, *Csf1r*^ΔFIRE/WT^ and *Fireko* mice were treated with CSF1-Fc on 4 successive days and analyzed on day 5. Prior to CSF1 administration, we assessed the body weight, fat mass and lean mass by NMR (Minispec) and found no significant difference between groups (**Figure S2A**), despite the reported role of CSF1R-dependent macrophages in adipose tissue development (Cox et al., 2021). CSF1-induced increases in liver weight and bone marrow and blood Ly6C^high^ monocyte numbers reported previously (Gow et al., 2014; Keshvari et al., 2024) were detected in both WT and heterozygous mice and entirely absent in *Fireko* mice (**Figure S2B-D).** Responses to CSF1-Fc that were not affected by the *Csf1r* mutation include thrombocytopenia and splenomegaly (**Figure S2E,F**) likely reflecting the fact that CSF1R expression in the spleen is not affected by the mutation (Rojo et al., 2019).

In summary, FIRE is required for expression of *Csf1r* in bone marrow progenitors and *Fireko* mice are CSF1 unresponsive. Monocytopoiesis does not absolutely require CSF1R expression in progenitor cells. Deletion of FIRE compromises but does not prevent expression of *Csf1r* in Ly6C^low^ blood monocytes.

### Csf1r expression in Fireko mice is inducible by CSF2

Consistent with absence of detectable CSF1R, bone marrow cells from *Fireko* mice did not respond to CSF1 in liquid culture as shown previously (Rojo et al., 2019). By contrast, cells from WT, *Csf1r*^ΔFIRE/WT^ and *Fireko* mice responded equally to granulocyte-macrophage colony-stimulating factor (GM-CSF, CSF2) to generate a spectrum of myeloid cells. In WT cultures, two major subpopulations were identified: F4/80^high^/CSF1R^high^/MHCII^low^ and F4/80^low^/CSF1R^low^/MHCII^high^ (**Figure 2A**). The latter population includes a mixture of classical dendritic cells and macrophage-like antigen presenting cells (Helft et al., 2015). Both populations were able to internalise pH Rodo-labelled *E.coli* bioparticles but the uptake measured as median fluorescence intensity was greater in MHCII^low^ cells (**Figure 2B**). Within the MHCII^low^ macrophage population, CSF1R expression in *Fireko* was detected at 10-15% of the WT level. Expression in *Csf1r*^ΔFIRE/+^ macrophages was intermediate between the two **(Figure 2A**). These observations confirm that CSF2 can promote the expression of *Csf1r* in *Fireko* bone marrow progenitors but does not entirely overcome the effect of the enhancer deletion.

**Figure 2.**
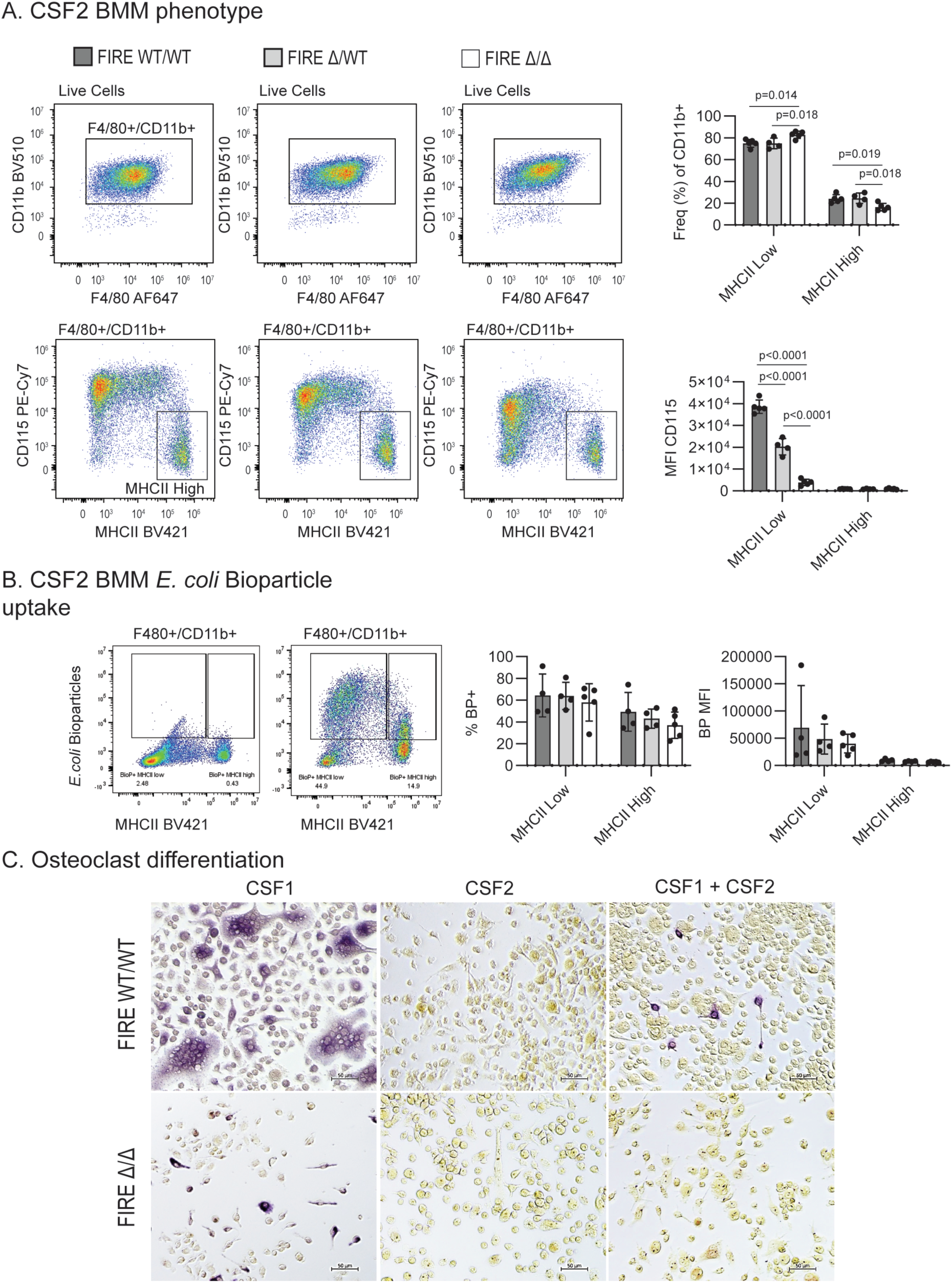
Regulation of CSF1R expression and differentiation in *Fireko* mice by CSF2. (A) Bone marrow-derived macrophages (BMM) from WT and *Fireko* mice were generated by culture in CSF2 for 7 days followed by staining and flow cytometry analysis. F4/80/CD11b^+^ cells were gated based on MHCII expression. The relative proportions of MHCII High and Low subsets and their CSF1R MFI are shown. Data show mean and standard deviation, 2-way ANOVA with Sidak’s multiple comparison test. (B) CSF2-derived BMM were incubated with pH Rodo *E.coli* bioparticles (BP) for 1 h followed by surface staining and analysis by flow cytometry for BP uptake (%) and median fluorescence intensity (MFI) in cell subsets. Data show mean and standard deviation. (C) Representative images of bone marrow from WT or *Fireko* mice cultured in CSF1, CSF2 or both for 3 days prior to the addition of RANKL and continued culture until day 7. Cells were stained for tartrate-resistant acid phosphatase (TRAP, purple)

Mice with mutations in *Csf1* or *Csf1r* lack osteoclasts (OCL) and are osteopetrotic (Dai et al., 2002) whereas *Fireko* mice have normal bone development. We next asked whether bone marrow cells from *Fireko* mice could generate OCL in response to stimulation with RANK ligand (Ovchinnikov et al., 2010). Bone marrow cells were cultured in CSF1, CSF2 or the combination for 3 days prior to addition of RANKL and continued culture until day 7. In WT bone marrow, addition of RANKL to cultures expanded in CSF1 led to the generation of multinucleated osteoclasts expressing tartrate-resistant acid phosphatase (TRAP). This response was absent in cells grown in CSF2 and prevented in the continued presence of CSF2 (**Figure 2C**). Hence CSF2 is unlikely to be responsible for the generation of OCL in the *Fireko*.

### Fireko mice respond to inflammation in the peritoneal cavity

The effect of CSF1/CSF1R inhibition on the generation of an inflammatory exudate in the peritoneal cavity has varied in previous studies which used different antibodies and treatment regimens (Louis et al., 2015; MacDonald et al., 2010). However, thioglycolate-elicited peritoneal macrophages (TEPM) are autocrine for CSF1R signaling (Irvine et al., 2006) and ChIP-seq analysis of TEPM revealed multiple active regulatory elements within the *Csf1r* locus in addition to FIRE (Fonseca et al., 2019). We therefore asked whether resident peritoneal macrophages, which are absent in *Fireko* mice, are required to initiate the response, and whether inflammatory recruitment could enable expression of CSF1R. Thioglycolate injection led to the loss of large F4/80^High^ peritoneal macrophages, the macrophage disappearance reaction (Barth et al., 1995), in WT mice as expected, and accumulation of F4/80^Low^ macrophages (derived from recruited monocytes) regardless of genotype (**Figure 3A,B**). Surface CD115 was detected on *Fireko* TEPM but at significantly lower levels than in WT and heterozygous mice (**Figure 3C**). Interestingly, granulocyte recruitment was reduced in the *Fireko* mice (**Figure 3B**), suggesting a role for resident macrophages in acute inflammation. We also analyzed the peritoneal macrophage populations in mice treated with CSF1-Fc (**Figure 3B,C**). CSF1-Fc treatment caused an increase in total yield of lavage cells and an increase in the % of large peritoneal macrophages within the lavage in WT mice. The peritoneal macrophage expansion in response to CSF1-Fc was significantly attenuated in the heterozygous mutant mice and the treatment did not overcome the macrophage deficiency of the *Fireko* mice.

**Figure 3.**
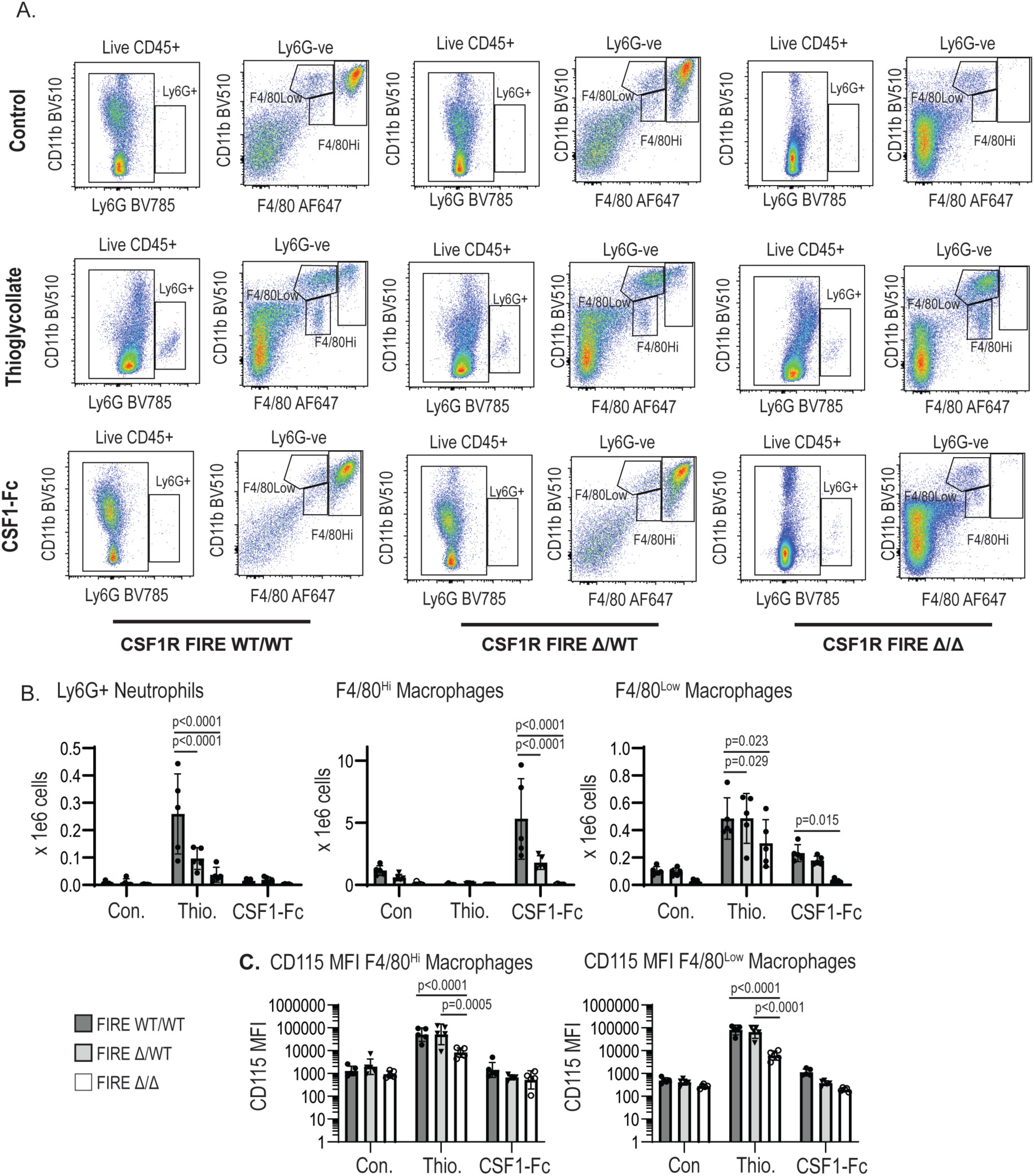
Analysis of the effect of homozygous and heterozygous ΔFIRE mutation on resident and elicited peritoneal macrophages. WT, *Csf1r*^ΔFIRE/+^ and *Fireko* mice on were administered 1 ml thioglycolate ip, or 4 x subcutaneous injections of 1 mg/kg CSF1-Fc or saline control. Peritoneal lavage was collected 5 days post thioglycolate or CSF1-Fc administration and analysed by flow cytometry. (A) Representative flow cytometry profiles of Ly6G^-^ populations defined by expression of CD11b and F4/80 following different treatment in each genotype. (B) Quantification of Ly6G^+^ neutrophils, F4/80^High^ and F4/80^Low^ macrophages in control and thioglycolate-elicited lavage. Each point is an individual animal. (C) Quantification of CD115 MFI on F4/80^High^ and F4/80^Low^ macrophages in control and thioglycolate-elicited lavage. Separate panels in (B) and (C) show quantification of total white blood cells, Ly6G^+^ neutrophils, F4/80^High^ and F4/80^Low^ macrophages in peritoneal lavage from control and CSF1-Fc treated mice. Each point is an individual animal of the indicated genotype.

### The conserved AP1 element in FIRE contributes to the rate of Csf1r transcription but is not essential for maintenance of tissue macrophages

The FIRE sequence includes a core AP1-ETS element conserved from reptiles to mammals (Hume et al., 2017). ChIP-seq analysis revealed binding of multiple AP1 family members to FIRE (Fonseca et al., 2019). The AP1 sequence contains the transcription start site of an anti-sense enhancer RNA that is induced by stimuli that repress *Csf1r* transcription (Sauter et al., 2013). To test the function of this element, we mutated the AP1 site in the mouse germ line to abolish AP1 binding. Comprehensive analysis of tissue macrophage populations in *Csf1r*^ΔAP1-FIRE/ΔAP1-FIRE^ mice revealed no loss of any of the populations missing in *Fireko* mice including microglia (**Figure S3A**). Peritoneal macrophage populations and their expression of surface CSF1R were unchanged compared to WT (**Figure S3B**). We next asked whether mutation impacted CSF1 responses *in vitro. Csf1r*^ΔAP1-FIRE/ΔAP1-FIRE^ bone marrow cells were significantly less responsive to CSF1 (**Figure 4A**). Bone marrow-derived macrophages (BMDM) were cultured in CSF1 to maximally down-regulate surface protein and *Csf1r* mRNA, then washed to remove the ligand, and reappearance of surface CSF1R (CD115 antigen) was measured with time. The *Csf1r*^ΔAP1-FIRE/ΔAP1-FIRE^ mutation reduced the rate of resynthesis of the receptor following CSF1 removal and compromised expression of *Csf1r* mRNA (**Figure 4B**). Surface CSF1R on BMDM is acutely down-regulated via ectodomain cleavage following protein kinase C (PKC) activation using phorbol myristate acetate (PMA) and by TLR4 activation following addition of lipopolysaccharide (LPS) (Sester et al., 1999). In the case of PMA, the effect is transient, due to degradation of PKC, and CSF1R reappears. The AP1 mutation also decreased the rate of reappearance of CSF1R in PMA-treated cells (**Figure 4B**). However, the mutation did not allow CSF1R protein, or *Csf1r* mRNA, to be re-expressed in the presence of LPS.

**Figure 4.**
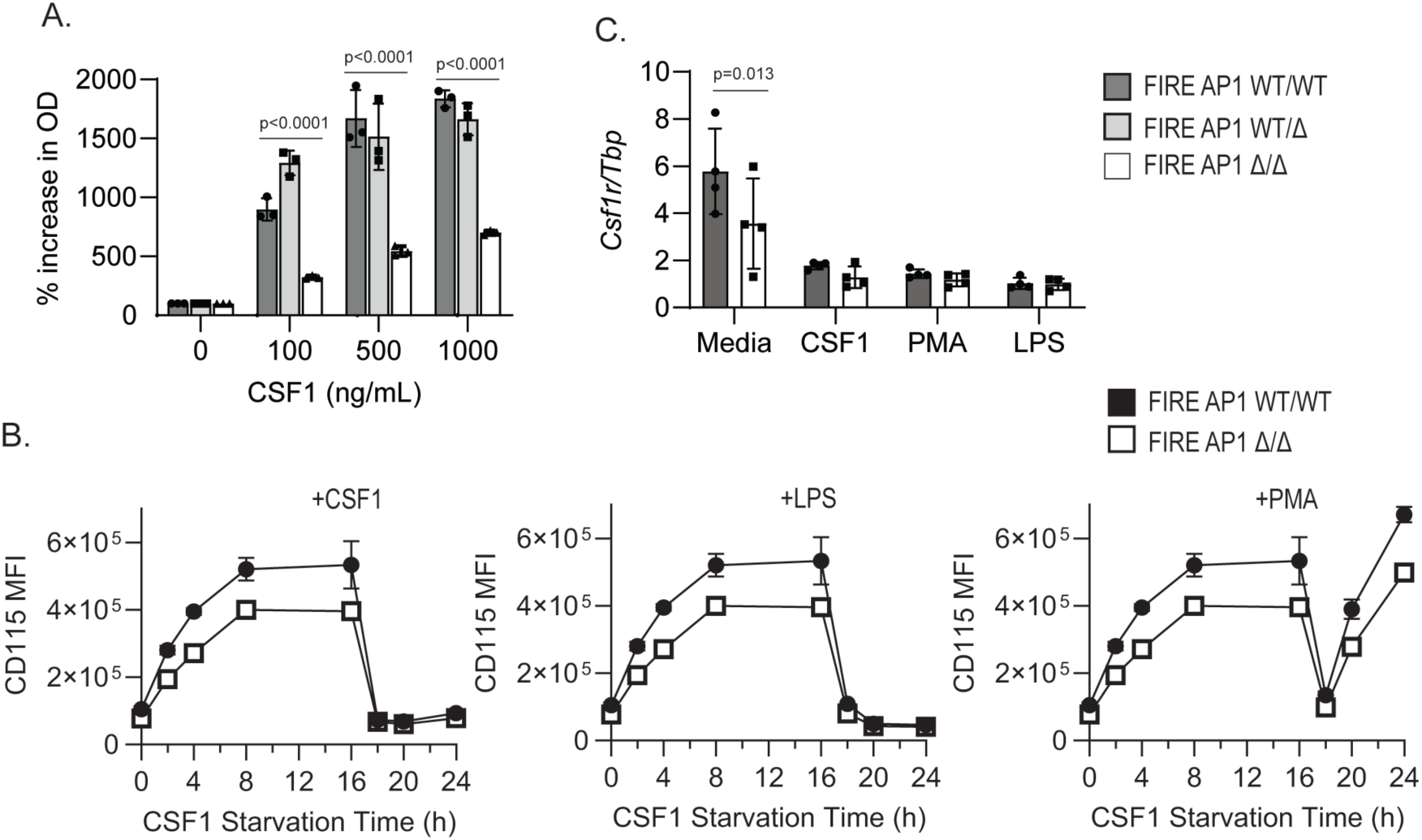
The *Csf1r*^ΔAP1-FIRE/ΔAP1-FIRE^ mutation regulates CSF1R expression. (A) Bone marrow cells from *Csf1r*^+/+^, *Csf1r*^+/ΔAP1^ or *Csf1r*^ΔAP1/ΔAP1^ mice were cultured for 7 days in recombinant human CSF1 at the indicated concentrations and metabolic activity was measured by resazurin assay. (B) CSF1-differentiated bone marrow macrophages were starved of CSF1 for 2, 4, 8, 16, and 24 hours, or starved of CSF1 for 16 hours followed by stimulation with CSF1, LPS or PMA for 2, 4, and 8 hours and surface expression of CSF1R (CD115) was analyzed by flow cytometry. MFI: median fluorescence intensity. (C) CSF1-differentiated bone marrow macrophages were starved of CSF1 for 16 h followed by stimulation with medium control, CSF1, PMA or LPS for 8 h. *Csf1r* mRNA expression was analyzed by qPCR. 2-way ANOVA with Sidak’s multiple comparisons test.

### Fireko mice lack defined subpopulations of tissue resident macrophages. Evidence of haploinsufficiency in heterozygous mutants

*Csf1r* mRNA and surface CD115 are each reduced by 50% in monocytes and macrophages in heterozygous *Csf1r*^ΔFIRE/+^ mice compared to WT (Rojo et al., 2019). Tissue resident macrophages, including microglia, undergo a CSF1-dependent expansion in the first weeks of postnatal life (Summers and Hume, 2017) that was delayed in mice with a heterozygous kinase dead *Csf1r* mutation (Stables et al., 2022). In view of this evidence and the apparent haploinsufficiency we speculated that *Csf1r*^ΔFIRE/WT^ mice might exhibit a similar delay. We found that microglial density was significantly reduced in *Csf1r*^ΔFIRE/WT^ mice compared to WT at 3 weeks of age (**Figure 5A,B**). By contrast to the reported microgliosis reported in *Csf1r^+/-^*mice on the C57BL/6J background (Chitu et al., 2015), there was no significant difference between WT and *Csf1r*^ΔFIRE/WT^ mice at 10 or 30 weeks (**Figure 5A,B**).

**Figure 5.**
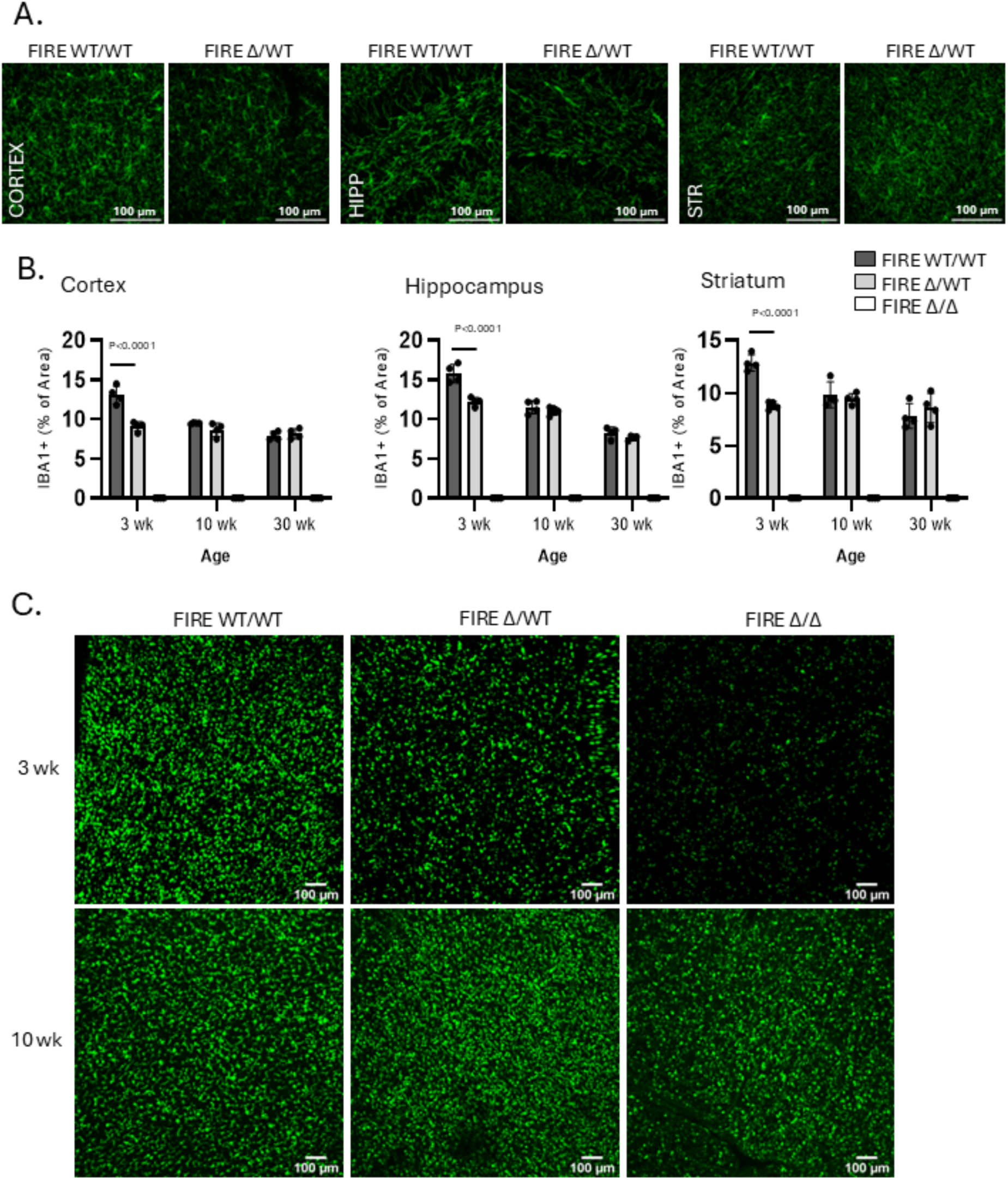
Depletion of microglia and lung macrophages in *Csf1r*^ΔFIRE/+^ mice. Brain sections from WT, *Csf1r*^ΔFIRE/+^ and *Fireko* mice at 3, 10 or 30 wks of age were stained for IBA1 in the cortex, hippocampus and striatum (A) Representative images from 3 wks. (B) Quantification of IBA1^+^ stained area by image analysis. Each point is an individual animal. Data show mean and standard deviation. 2 way ANOVA with Tukey’s multiple comparisons test. (C) Representative whole mount images of *Csf1r-*EGFP in WT, *Csf1r*^ΔFIRE/+^ and *Fireko* mice at 3 and 10 weeks.

Subsequent to the original description of the *Fireko* mice (Rojo et al., 2019), others described partial macrophage deficiency in salivary gland (McKendrick et al., 2023) and omentum (Louwe et al., 2022). To fully evaluate the extent of macrophage deficiency in these mice we used whole mount imaging of the *Csf1r*-EGFP reporter (Sasmono et al., 2003). *Csf1*^op/op^ mice have a deficiency in alveolar macrophages at 3 weeks that resolves with age (Shibata et al., 2001). The abundance of SiglecF-positive alveolar macrophages was previously found to be unaffected in adult *Fireko* mice (Rojo et al., 2019). A separate population of interstitial macrophages is poorly recovered following tissue disaggregation (Hume et al., 2023). In WT embryonic lung, *Csf1r-*EGFP is expressed by two morphologically distinct populations of interstitial macrophages, distinguished by mutually exclusive expression of MAC2 (*Lgals3*) and F4/80 respectively. *Csf1r-*EGFP^+^ interstitial (F4/80^+^) and alveolar (MAC2^+^) macrophages expand during the first 3 weeks of postnatal life (Tan and Krasnow, 2016). At 3 weeks of age, *Fireko* mice were deficient in *Csf1r-*EGFP^+^ cells in the lung, heterozygous mutant mice appeared intermediate in phenotype (**Figure 5C**). By 10 weeks of age there was no longer a clear distinction between genotypes, consistent with the previous analysis of adult *Fireko* mice(Rojo et al., 2019).

The testis contains two distinct macrophage populations with distinct ontogeny, interstitial macrophages that interact with Leydig cells and regulate steroidogenesis and peritubular macrophages that interact with Sertoli cells (Gu et al., 2023). Both are readily imaged in adult testis using the *Csf1r-*EGFP reporter and the stellate peritubular population is selectively depleted in *Fireko* mice (**Figure S4A**). The loss of peritubular cells was confirmed by localisation of F4/80 (**Figure S4A**). Langerhans cells (LC), the macrophages of the epidermis, are also derived from embryonic progenitors and expand rapidly in the postnatal period (Mass et al., 2023). We confirmed that LC were depleted in the ear of adult mice as reported previously (**Figure S4B**) (Rojo et al., 2019). At 3 weeks of age, *Fireko* mice were devoid of detectable LC in the more stratified epidermis of the footpad (not shown) but this deficiency was fully resolved by 10 weeks (**Figure S4B**).

The macrophages of the pancreas were not previously examined in the *Fireko* mice. Three distinct populations have been identified, two interstitial macrophages within the exocrine pancreas that differ in self-renewal capacity and a population within the islets (Calderon et al., 2015). Based upon detection of *Csf1r-EGFP* in whole mounts, interstitial macrophages were reduced in *Fireko* mice at 10 weeks but partly replenished at 30 weeks (**Figure S4C**). *Fireko* mice lack cardiac muscle macrophages on the inbred background (see below). The abundance of *Csf1r-EGFP* macrophages was also greatly reduced in skeletal muscle in the diaphragm, which can be readily imaged in whole mounts (**Figure S4D**). In adipose tissue at 10 weeks, we noted an apparent reduction in stellate interstitial *Csf1r-*EGFP-positive macrophages in epididymal fat pads (**Figure S4E**).

The liver contains a subcapsular network of CSF1-dependent macrophages that can be visualized with the *Csf1r* reporter transgene (Sierro et al., 2017). These cells are absent in the *Fireko* liver, whereas the underlying F4/80^+^ Kupffer cells are retained (**Figure S5A**). By contrast, the CD169^+^ marginal zone metallophils of the spleen, which are also CSF1/CSF1R dependent (Stables et al., 2024; Witmer-Pack et al., 1993) were unaffected in *Fireko* mice (**Figure S5B**).

**Figure S6** contains additional representative immunohistochemistry images and quantitation of tissue resident macrophages in adult mice from the congenic C57BL/6J *Fireko* line. F4/80^+^ macrophages were present in white and brown adipose tissue in both male and female WT and *Fireko* mice (Figure 6A). By contrast to reported depletion of F4/80^+^ macrophages in salivary gland on the original background (McKendrick et al., 2023) these cells were partly retained (**Figure S6B**). Interstitial macrophages in the pancreas were not readily detected with F4/80 so IBA1 was used as a marker. This revealed a selective reduction in interstitial macrophages in the *Fireko*, consistent with EGFP detection, whereas a distinct population on the surface of lobules was increased (**Figure S6C**). F4/80 was detected on macrophages within islets of Langerhans, and this population was absent in *Fireko* mice (**Figure S6D**). In other endocrine tissues, the abundant F4/80^+^ macrophage population of the adrenal cortex and the microglia-like population of the medulla (Hume et al., 1984) were both absent in the *Fireko* (**Figure S6E**). In the *Fireko* ovary, interstitial F4/80^+^ macrophages were reduced and the recruited macrophages that populate the WT corpus luteum (Hume et al., 1984) were absent (**Figure S6F**). Finally in pituitary, the microglial-like cells of the posterior lobe were absent in the *Fireko* and those of the anterior lobe were reduced, but a border-associated macrophage population was unaffected (**Figure S6G**).

**Figure 6.**
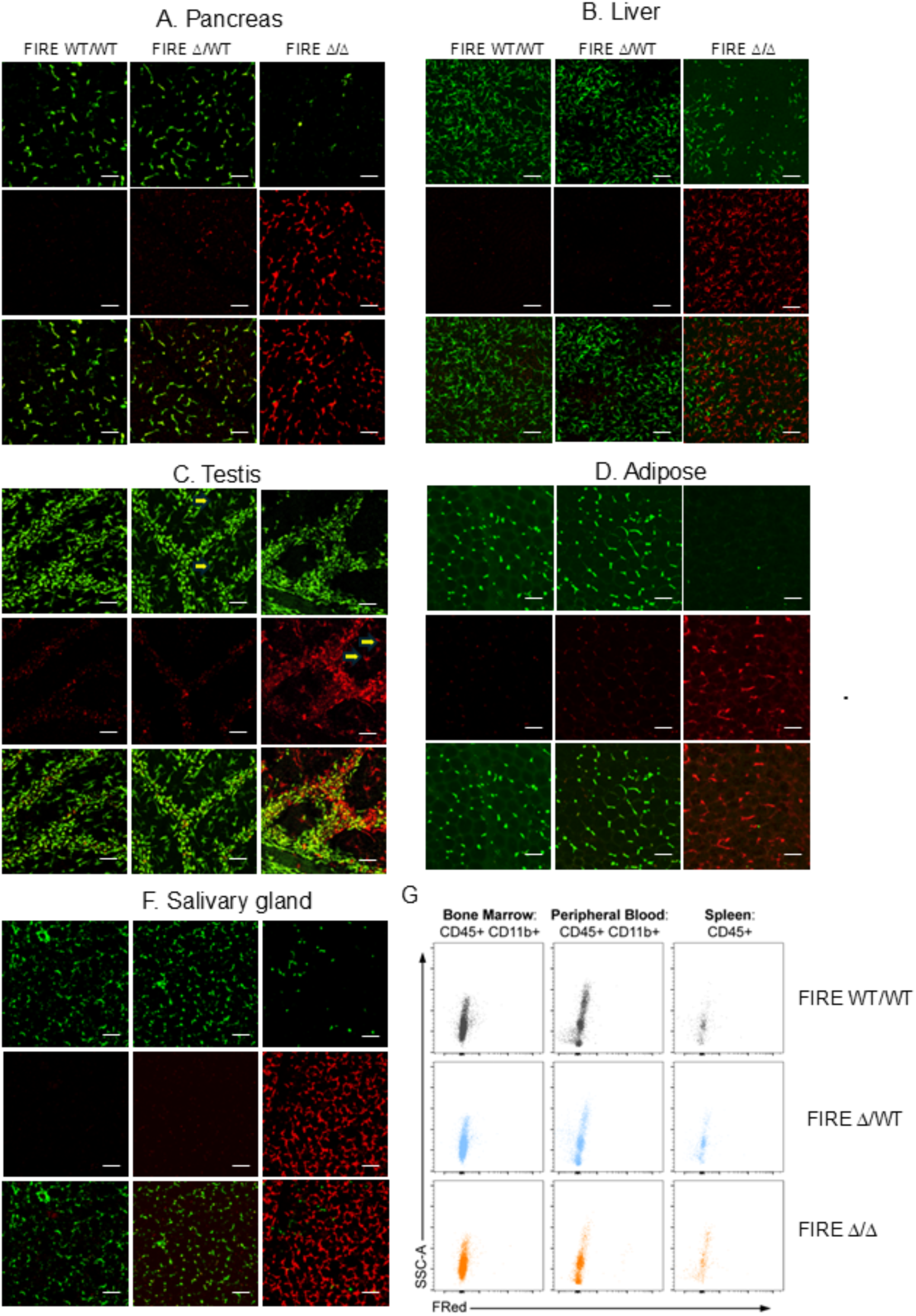
Donor cell population of tissue macrophages in *Fireko* mice following transfer of WT bone marrow. *Csf1r-*EGFP^+^ WT, *Csf1r*^ΔFIRE/+^ and *Fireko* mice were injected i.p. at weaning with WT *Csf1r-*FRed donor cells and chimerism was assessed by whole mount imaging 12 wks post transfer. Images of selected tissues in Panels A-F are representative of at least 5 BMT recipients analyzed. Scale bars=50μm. Panel G shows typical flow cytometry profiles of bone marrow, blood and spleen cells demonstrating the absence of detectable FRed^+^ donor cells following bone marrow transfer.

### Renal macrophage deficiency in Fireko mice does not impact basal kidney function or the development of chronic kidney disease

To analyse the impact of congenital macrophage deficiency on organ function we focused on the kidney. In terms of gene expression profile, resident kidney macrophages share some features with microglia (Summers et al., 2020). Adult *Fireko* mice were previously shown to lack F4/80^high^ kidney macrophages by flow cytometry analysis (Rojo et al., 2019). IBA1^+^ immunostaining highlighted the reduction in macrophages in cortex and medulla in both *Fireko* and heterozygous mutant mice (*Csf1r*^ΔFIRE/WT^) compared to WT (**Figure S7A,B**). By contrast to other organs, the deficiency in heterozygous mice did not resolve with age. Localization of F4/80 confirmed the complete absence of the abundant F4/80^high^ renal macrophage populations (Hume and Gordon, 1983) in *Fireko* mice (**Figure S7C).** Residual IBA1^+^ cells in the *Fireko* kidney medulla are likely the retained F4/80^low^ monocytes and monocyte-derived cells detected by flow cytometry (Rojo et al., 2019). *Csf1rko* in rats is associated with impaired postnatal kidney development including severe renal medullary hypoplasia (Keshvari et al., 2021). Figure S7A,B includes localization of the vascular marker CD31 in 3 and 10 week old WT and *Fireko* mice, highlighting glomeruli in the cortex and major vessels in the medulla. No gross anatomical differences were evident in the mutant mice. Whereas *Adgre1* mRNA was depleted in *Fireko* mice consistent with loss of detectable F4/80, the apparently normal histology and function of *Fireko* kidneys (see below) was supported by unchanged detection of mRNA encoding markers of podocytes (*Nphs2*), proximal tubules (*Slc22a13*) distal tubules (*Umod*) or collecting ducts (*Aqp2, Scnn1b*) in total kidney mRNA (**Figure S7D**).

To analyze the function of resident kidney macrophages in response to epithelial injury we exposed mice to dietary adenine. Adenine metabolism in rodents leads to the production of 2,8-dihydroxyadenine crystals, which accumulate in the renal tubules, causing mechanical obstruction, tubular injury, and inflammation, which is typically more severe in male mice (Diwan et al., 2018; Makhloufi et al., 2020; Miguel et al., 2021). Mice were administered normal chow or a diet containing 0.2% adenine for 4 weeks. Macrophages have been implicated in kidney development, including the renal vasculature (Alikhan et al., 2011; Munro et al., 2019; Rae et al., 2007). Despite profound macrophage deficiency the kidneys of control adult WT and *Fireko* mice were of similar size (**Figure S8B**) and were not distinguished based upon evaluation of H&E, periodic-acid-Schiff (PAS) or Sirius-red stained sections by a veterinary pathologist (RA) blinded to genotype (**Figure S8C**). Despite the paucity of F4/80^high^ macrophages in the *Fireko* kidneys at steady state, adenine injury promoted massive interstitial accumulation of F4/80^+^ macrophages in cortex and medulla in both WT and *Fireko* mice (**Figure S8C**). The *Fireko* had no effect on markers of renal function, serum creatinine or urea, in control chow fed male or female mice (**Figure S8E**). All adenine treatment groups lost weight but there was no difference between genotypes in body weight or kidney weight (**Figure S8A,B**). Kidneys from adenine-treated mice exhibited severe tubular pathology (atrophy, adenine crystal and cast/debris deposition), milder glomerular pathology (Bowman’s capsule thickening) and fibrosis, associated with increased circulating creatinine and urea (**Figure S8D**). Disease severity was similar between genotypes histologically but circulating creatinine and urea levels were significantly elevated in male *Fireko* mice compared to WT (**Figure S8E**).

### Cardiac macrophage deficiency in Fireko mice does not affect steady state heart function

The abundant tissue macrophages in the heart have been ascribed essential functions in electrical conductivity and metabolic homeostasis, with conditional depletion of macrophages leading to changes in cardiac hemodynamics and ventricular dysfunction within 3 weeks of treatment (Hulsmans et al., 2017; Nicolas-Avila et al., 2020). We localized cardiac resident macrophages using IBA1 in combination with CD169 and CD206 which have been reported to define subpopulations of cardiac macrophages (Wong et al., 2021). Those populations were depleted in the *Fireko* (Rojo et al., 2019). As observed in the kidney, the resident cardiac macrophages were also deficient in older *Csf1r*^ΔFIRE/+^ mice (**Figure S9A).** To determine the effect of cardiac macrophage deficiency we examined cardiac function in 13-19 week old animals by cardiac ultrasound (Nicolas-Avila et al., 2020). As shown in **Figure S9B**, none of the measured parameters was significantly different between wild-type and *Fireko* mice.

### Vacant tissue macrophage niches are populated by donor cells following adoptive transfer of WT bone marrow cells

Taking advantage of the inbred genetic background, WT, *Csf1r*^ΔFIRE/+^ and *Fireko* mice with the *Csf1r-*EGFP transgene were injected intraperitoneally with WT bone marrow cells (BMT) from *Csf1r-*FusionRed (FRed^+^) donors at weaning (3 weeks). Chimerism was assessed 12 weeks post BMT. In the periphery, whole mount imaging revealed the presence of donor FRed^+^ macrophages within the vacant territories identified in the *Fireko* recipients. Selected tissues are shown in Figure 6 and **Figure S10**. Surprisingly, although adipose, pancreas, ovary, diaphragm and salivary gland in adult *Fireko* mice retained populations of F4/80^+^/IBA1^+^ macrophages, in the BMT recipients macrophages with similar density and stellate morphology were entirely of donor (FRed^+^) origin. In the testis, we observed replacement of peritubular macrophages as well as foci of FRed^+^ interstitial macrophages (Figure 6C).

In the liver donor cells were detected amongst the capsular population, whereas underlying Kupffer cells remained of recipient origin (EGFP^+^, Figure 6B). Similarly, in the lung, donor cells populated the stellate interstitial population whereas alveolar macrophages identified by more rounded morphology and location within the airways remained EGFP^+^. We also detected donor cells on the surface of the spleen whereas the red pulp macrophages remained EGFP^+^ (**Figure S10**). Importantly, no FRed+ cells were detected by flow cytometry analysis of CD11b^+^ myeloid cells in bone marrow and blood or spleen cells of transplanted *Fireko* mice (Figure 6G).

The CD11b^high^ large peritoneal macrophage population, which is absent in *Fireko* mice, was populated by FRed^+^ donor cells (Figure 7A). No donor cells were detected in any organ in wild type recipients (Figure 6). However, in heterozygous recipients partial chimerism was detected in the peritoneum (Figure 7A) and in organs with direct access to the peritoneal cavity (diaphragm, adipose, pancreas, liver capsule (**Figure S10B**)). In the *Fireko* kidney, F4/80^high^ resident macrophages are selectively depleted whereas F4/80^low^ populations are retained (Rojo et al., 2019). Each of these populations express FRed in the donor, but in BMT recipients only the F4/80^high^ macrophages were of donor origin (**Figure S10C).**

**Figure 7.**
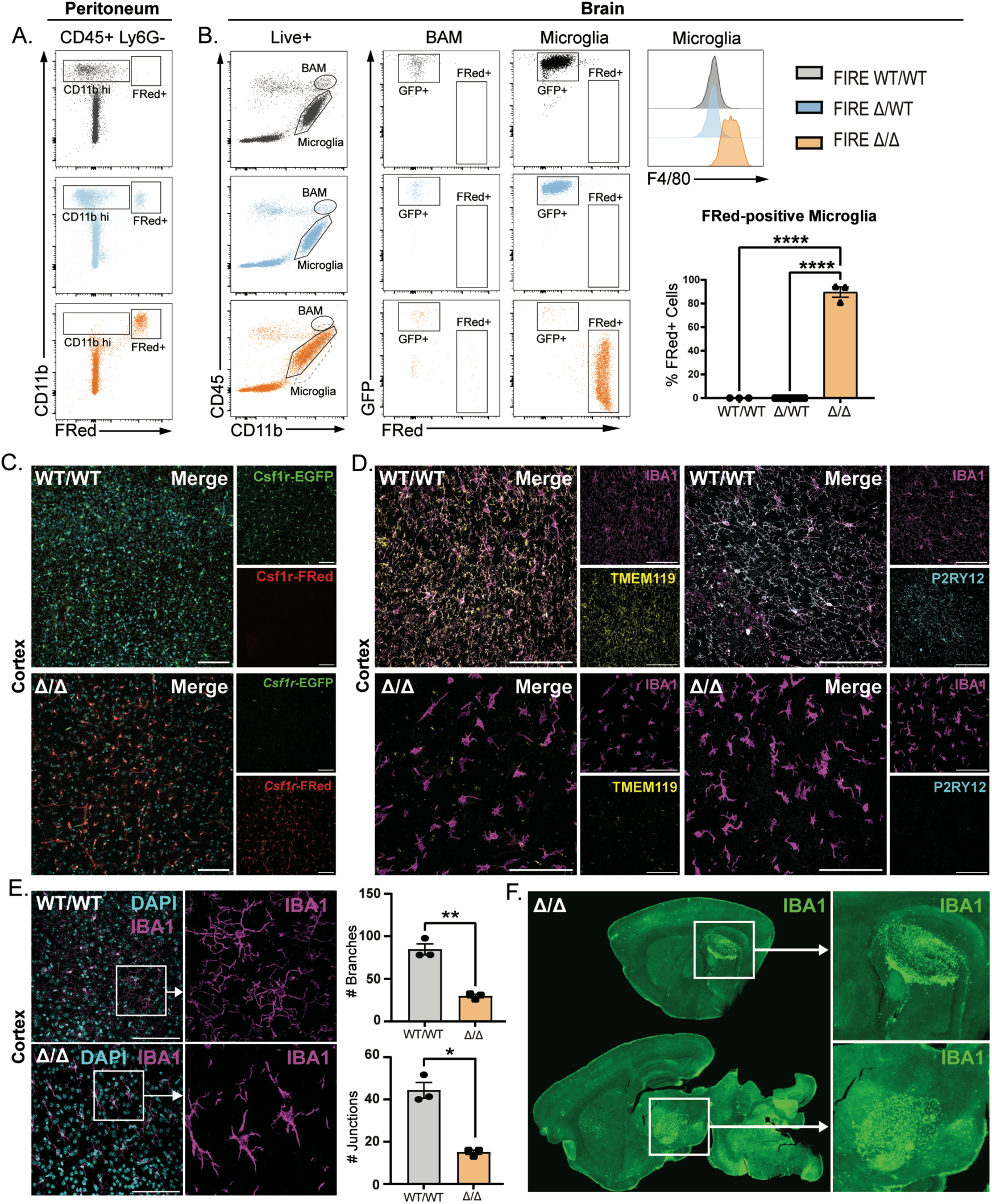
Repopulation of the brain of *Fireko* mice with WT donor cells following bone marrow transfer (BMT). *Fireko*/*Csf1r-*EFGP mice were injected intraperitoneally (IP) with *Csf1r*-FusionRed (Fred) donor bone marrow cells at weaning (3 weeks), and engraftment was analyzed 12 weeks post-transfer. (A) Representative flow cytometry profiles of peritoneal cells harvested from WT (+/+), *Csf1r*^ΔFIRE/+^ (Δ/+) and *Fireko* (Δ/Δ) recipients. (B) Representative flow cytometry profiles of brain myeloid cells harvested from the same cohort. The dotted line in the CD45/CD11b profile highlights the shift in location of the microglial gate from WT to *Fireko* BMT recipients. Proportions of FRed^+^ microglia are quantified on right. Graph shows mean ± SEM from at least three mice per genotype, and p values were determined using ordinary one-way ANOVA followed by Tukey’s multiple comparison test. ****, p < 0.0001. Histogram panel shows F4/80 staining on microglial cells isolated from BMT recipients. (C) Representative images of sections of brain cortex from WT (+/+) and *Fireko* (Δ/Δ) BMT recipients show the repopulation of the cortex with donor (FRed^+^ cells). Scale bar = 100 µm. (D) Representative images of sections of brain cortex from WT (+/+) and *Fireko* (Δ/Δ) BMT recipients stained for IBA1 (magenta), TMEM119 (yellow), and P2RY12 (teal). Scale bars = 100 µm. (E) Representative images of sections of brain cortex stained with IBA1 (magenta) and co-stained with DAPI. Highlighted regions (white box) are shown at higher resolution to the right to highlight the morphological difference between WT microglia and microglia repopulated in *Fireko* mice. Microglial ramification was quantified in ImageJ. Graphs show mean ± SEM of three mice per genotype, and p values were determined using an unpaired t-test with Welch’s correction. *, p < 0.05; **, p < 0.01. Scale bars = 100 µm. (F) *Fireko* (Δ/Δ) recipients were sacrificed at 3 weeks post BMT. Serial free-floating sections were stained to detect repopulation with IBA1^+^ cells. Every section examined contained multiple foci of IBA1^+^ cells, panels show typical examples.

The brain is of particular importance because of the neuropathology associated with *CSF1R* mutations in humans (Chitu et al., 2022) and the potential for treatment by microglial replacement. The brains of *Fireko* mice were populated with FRed^+^ donor-derived cells 12 weeks post-BMT (Figure 7B**,C**). No residual *Csf1r*-EGFP^+^ cells were detected in the brain parenchyma of BMT recipients, indicating that recipient BAMs were also replaced. No FRed^+^ donor cells were detected in the brains of WT or heterozygous BMT recipients (Figure 7B). Previous evidence indicated that bone marrow-derived microglia-like cells do not fully recapitulate homeostatic gene expression (Bastos et al., 2025; Bennett et al., 2018; Carter-Cusack et al., 2024; Cronk et al., 2018; Lund et al., 2018; Shemer et al., 2018; Shibuya et al., 2022). In BMT recipients, FRed^+^ donor-derived cells had relatively increased CD45 and F4/80 expression compared to WT microglia (Figure 7B). Donor-derived cells were uniformly strongly FRed^+^ and achieved a similar density and regular distribution to WT microglia. However, they did not replicate the extensive ramification characteristic of normal microglia (Figure 7C) and expression of homeostatic microglia markers P2RY12 and TMEM119 was barely detectable in the donor-derived cells (Figure 7D). The difference in morphology between resident and BM-derived cells in the cortex is shown in more detail by IBA1 staining and quantitative image analysis in Figure 7E. To gain insight into the mechanism of repopulation of the microglial niche, we examined brains at three- and five-weeks post BMT. Within individual brains we observed multiple large clusters of IBA1^+^ cells with a concentrated “wave front” similar to the pattern detected following intracerebral microglial transplantation (Chadarevian et al., 2024) (Figure 7F).

## Discussion

### Selective loss of resident tissue macrophage populations in Fireko mice

The *Csf1r*-EGFP transgene (Sasmono et al., 2003) is expressed by all myeloid cells and some B cells. In most tissues it is co-expressed with F4/80 and provides a convenient marker to enable visualization of tissue mononuclear phagocytes. It was used previously to demonstrate the depletion of resident tissue macrophages following extended treatment with anti-CSF1R antibody (MacDonald et al., 2010). In the current study we combined the *Csf1r-*EGFP reporter with the *Csf1r*^ΔFIRE^ allele on the congenic C57BL6/J background to extend the previous analysis of *Fireko* mice by flow cytometry on disaggregated tissues (Rojo et al., 2019). In the kidney, we demonstrated the absence of F4/80^high^ resident macrophages in the cortex and the more abundant population lining the tubules in the medulla (Hume and Gordon, 1983). Based upon the loss of both *Csf1r-*EGFP and F4/80, several additional tissue macrophage populations were found to be depleted in the tissues of *Fireko* mice including subpopulations in liver, lung, skeletal muscle, testis, ovary, pituitary, adrenal and pancreas. The testis contains two distinct macrophage populations occupying interstitial and peritubular locations (DeFalco et al., 2015; Gu et al., 2023; Gu et al., 2022; Mossadegh-Keller et al., 2017).The peritubular macrophages have been considered CSF1R-negative, based upon binding of CD115 to isolated cells (DeFalco et al., 2015) but they were readily detected with *Csf1r-*EGFP and also in *Csf1r*-FRed mice (Grabert et al., 2020). The CSF1R^low^ population was selectively lost in the mutant mice confirming that the peritubular cells occupy a separate niche and the two testis populations are regulated separately. The lung contains an interstitial macrophage population that is enriched in the periphery of the organ (Tan and Krasnow, 2016) and is readily imaged in whole mounts. This population was selectively depleted in the juvenile *Fireko* mice. We speculate that the perinatal lethality associated with *Csf1r* mutations on the C57BL/6J background (Chitu and Stanley, 2017) may be related to neonatal lung macrophage deficiency (Abe et al., 2025). The original description of the *Fireko* mice described the loss of epidermal Langerhans cells (LC) in the ear skin. In thicker stratified squamous epithelia, LC spread in three dimensions through multiple layers of epidermal cells. We found that sustained loss of LC in the *Fireko* mice was specific to the epidermis of the ear where LC are located in the basal cell layer of individual squame piles (Hume et al., 1983). This may relate in part to ontogeny. Mass *et al*. (Mass et al., 2023) reviewed evidence of monocytic contribution to generation and maintenance of LC and dermal macrophages in different locations.

### Heterozygous phenotype in Csf1r^ΔFIRE/+^ mice indicates quantitative CSF1R signaling

The rapid postnatal expansion of resident macrophage population in all mouse tissues is detectable in total tissue transcriptomic profiles (Summers and Hume, 2017). This expansion is CSF1/CSF1R dependent(Stables et al., 2022). Here we found that heterozygous *Csf1r*^ΔFIRE^ mutation was sufficient to compromise postnatal expansion of macrophage populations in tissues affected by the homozygous mutation (brain, kidney, heart). These findings indicate that receptor availability is limiting for response. This in turn depends on continuous replacement of CSF1R on the surface as ligand-bound receptor is internalized and degraded (Stanley and Chitu, 2014). In essence, the response to CSF1 is determined not by ligand concentration but by the kinetics of receptor synthesis and trafficking. The highly-conserved AP1 element of FIRE contributes to the kinetics of receptor re-expression paralleling the requirement for a similar conserved ETS/AP1 element in an enhancer of the CSF1R target gene, *Plau* (Fowles et al., 1998). The homozygous AP1 mutation led to a reduction in the rate of reappearance of CSF1R following growth factor removal and reduced CSF1-dependent proliferation. Transient macrophage depletion seen in juvenile heterozygotes cannot be sustained because of the simple homeostatic feedback mechanism; reduced numbers of CSF1R^+^ macrophages consume less of the available local growth factor (Sehgal et al., 2021). The continued depletion of macrophages in kidney and heart in heterozygous mice may indicate that they depend on circulating rather than local CSF1. In the context of CSF1R-related leukoencephalopathy, the results do indicate a possible impact of haploinsufficiency that could interact with genetic background or an environmental trigger to initiate disease (Hume, 2025).

### Mechanisms contributing to niche-specific loss of tissue resident macrophages

In locations where the F4/80^high^ resident population is depleted in *Fireko* mice, the residual macrophage populations are predominantly F4/80^low^, monocyte-derived cells (McKendrick et al., 2023; Rojo et al., 2019; Weinberger et al., 2024). The prevailing view based upon fate-mapping studies is that most tissue macrophage populations in C57Bl/6J mice are established during embryonic development and maintained by self-renewal independent of input from bone marrow progenitors (Hume, 2023). The kidney, which like the brain is macrophage deficient in *Fireko* mice, is one clear exception with embryonic macrophage populations being replaced by bone marrow derived cells in the first few weeks of life (Salei et al., 2020). *Fireko* mice are macrophage-deficient in the embryo (Munro et al., 2020). Even those organs that retain macrophages in adult *Fireko* mice, such as liver, lung, intestine and spleen, were macrophage-deficient in the embryo and remain partly macrophage deficient at 3 weeks of age. Delayed development of resident macrophages in *Fireko* mice may explain why donor cells transferred at weaning were able to populate organs that were not macrophage-deficient in adults (Figure 6). Interestingly, macrophages of the intestinal lamina propria are believed to turn over continuously from blood monocytes (Bain et al., 2014) and this population is depleted by anti-CSF1R treatment (MacDonald et al., 2010). Yet, expression profiling of the intestine of WT and *Fireko* mice revealed no significant difference in detection of markers of monocytes (*Ccr2, Cx3cr1, Cd14*) or long-lived resident macrophages (*Csf1r, Adgre1, Folr2, Timd4*) (Rojo et al., 2019).

*Fireko* mice lack *Csf1r* expression in bone marrow progenitors and monocytes and are unresponsive to CSF1 *in vitro* or *in vivo* but they are not monocyte-deficient in marrow or blood. This observation is consistent with previous evidence that a blocking anti-CSF1R antibody does not deplete monoblasts in the marrow (Sudo et al., 1995) or blood monocytes (Louis et al., 2015; MacDonald et al., 2010). Blood monocyte subsets defined by markers such as Ly6C, CCR2 and CX3CR1 have been considered a developmental series that depends upon CSF1R signaling (Guilliams et al., 2018; Thierry et al., 2024). Ly6C^high^ classical monocytes may regulate the survival of their Ly6C^low^ progeny by acting as a sink for CSF1 (Yona et al., 2013). The loss of Ly6C^high^ monocytes in CCR2^-/-^ or anti-CCR2-treated mice led to decreased surface CSF1R and increased circulating half-life of Ly6C^low^ monocytes. Treatment with anti-CSF1R antibody in mice leads to the loss of non-classical monocytes and accumulation of classical monocytes (Louis et al., 2015; MacDonald et al., 2010). However, despite the almost complete loss of CSF1R in progenitors and monocytes in bone marrow, the relative abundance of circulating Ly6C^low^ monocytes was not reduced in *Fireko* mice. The *Csf1r-*FRed reporter is expressed around 5-fold higher in Ly6C^low^ compared to Ly6C^high^ monocytes (Grabert et al., 2020). The Ly6C^low^ monocytes had detectable, albeit reduced, CSF1R expression, when incubated without CSF1 *in vitro*. Both decreased CSF1R and increased abundance of Ly6C^low^ monocytes are likely due to the absence of CSF1R on Ly6C^high^ monocytes effectively removing the competition for endothelial CSF1 (Thierry et al., 2024). Consistent with this view, treatment of the *Fireko* mice with anti-CSF1R antibody depleted nonclassical monocytes confirming that they remain dependent on CSF1 (Gallerand et al., 2025).

The macrophage populations of the major organs that retain tissue macrophages in *Fireko* mice, liver, spleen, intestine and lung, are also partly retained in *Csf1r* knockout mice (Dai et al., 2002); preprint;(Jacquelin et al., 2025) and rats (Carter-Cusack et al., 2024; Keshvari et al., 2021) suggesting that these organs provide factors that can maintain survival and induce CSF1R expression in infiltrating monocytes. CSF2 in lung and intestine is one such factor (Kvedaraite et al., 2024; van de Laar et al., 2016; Varol et al., 2009); intestinal CSF2 in portal blood could also contribute to retention of liver Kupffer cells. However, although CSF2 promoted proliferation, differentiation and detectable CSF1R in bone marrow cells from *Fireko* mice, it did not restore WT levels of expression and actually prevented osteoclast differentiation. In bone, *Fireko* mice are not osteoclast-deficient (Rojo et al., 2019). The analysis in Figure 1G indicates that F4/80^+^ hematopoietic island macrophages are also retained and can express CSF1R, by contrast to Ly6C^high^ marrow monocytes. Two recent studies have provided evidence that conditional deletion of *Csf1* expression in marrow adipogenic stromal cells leads to loss of osteoclasts and resident macrophages without affecting monocytes (Inoue et al., 2023; Zhong et al., 2023).

FLT3 ligand (FLT3LG) has been identified as the factor that sustains osteoclast progenitors in *Csf1*^op/op^ mice and promotes age-dependent recovery (Lean et al., 2001). The relative importance of FLT3LG in monocytopoiesis and tissue macrophage development differs between humans and C57BL/6J mice (Momenilandi et al., 2024) but like CSF1, FLT3LG does cause a profound monocytosis when administered to mice (Brasel et al., 1996). In mice, analysis of double mutants indicates overlapping and partly compensatory functions of CSF1R and FLT3 in conventional dendritic cell differentiation (Durai et al., 2018; Percin et al., 2018). Their interactions in mouse monocyte-macrophage development remain to be tested.

In those organs that lack resident tissue macrophages in the *Fireko* mice, the monocyte-derived populations, which turn over relatively rapidly in WT mice (e.g. as shown in the kidney (Salei et al., 2020)) are generally retained. These monocyte-derived cells should have the capacity to differentiate to generate resident macrophages and fill the vacant niche (Bain et al., 2020; Louwe et al., 2022) but this process requires the continued expression of CSF1R. In overview, we suggest that monocyte homeostasis; marrow production and release, circulating half-life, extravasation, recruitment in response to inflammatory stimulus and tissue turnover are only partly disrupted in *Fireko* mice despite the loss of CSF1R in progenitors.

### Transcriptional regulation of Csf1r

The FIRE enhancer is clearly required for maximal expression of *Csf1r* in progenitors and blood monocytes and the requirement remains in bone marrow-derived macrophages grown in CSF2 and in recruited inflammatory macrophages. ChIP-seq analysis has revealed that FIRE is bound by numerous myeloid-specific transcription factors including IRF8, PU.1, CEBPA and MAFB (Kim et al., 2024; Rojo et al., 2017; Vanneste et al., 2026). In the current study we examined the effect of mutating the most highly-conserved element within FIRE, the compound AP1/ETS element. The disruption of this site impacted the kinetics of re-expression of *Csf1r* mRNA and protein following down-regulation and reduced the proliferative response to CSF1 but did not phenocopy the homozygous FIRE deletion. However, FIRE is not the only regulatory element identified by chromatin analysis within the *Csf1r* locus (Hoeksema et al., 2021; Link et al., 2018; Rojo et al., 2017). ATAC-seq analysis revealed regions of open chromatin that were specific to isolated macrophage populations that likely explain tissue-specific expression of *Csf1r* (Rojo et al., 2017). In the lung, peaks of open chromatin detected by ATAC-seq aside from FIRE distinguish AM, interstitial macrophages and monocytes (Sajti et al., 2020). In the liver, NOTCH2 and BMP signaling contributed to the differentiation of KC from blood monocytes recruited to the liver following KC ablation (Bonnardel et al., 2019; Sakai et al., 2019). Sakai *et al*. (Sakai et al., 2019) generated ATAC-seq data and ChIP-seq for key transcription factors, RBPJ, LXR (NR1H3) and SMAD4 for the monocyte-KC differentiation series. UCSC browser tracks from this study identify multiple KC-specific *Csf1r* enhancers in addition to FIRE, several of which bind the three regulators in both recruited monocytes and KC. These data may provide one explanation for the redundancy of FIRE in monocyte-derived KC.

### The absence of resident tissue macrophages in specific organs does not compromise development and homeostasis

Macrophages have been attributed numerous functions in embryonic and postnatal development and physiology (Mass et al., 2023; Matsudaira and Prinz, 2022; McNamara et al., 2023; Zhao et al., 2024). Thus far analysis of the *Fireko* mice has not revealed any essential physiological function of resident macrophages. By contrast to the infertility of *Csf1* and *Csf1r* mutant mice (Dai et al., 2002), and despite loss of macrophages in the gonads, *Fireko* mice are fertile. Renal medullary macrophages were recently shown to have a role in monitoring urine contents to prevent accumulation of sedimentary particles (He et al., 2024) but we found no histological evidence of tubular blockage or renal dysfunction in their absence. We also did not replicate the reported impact of macrophage depletion on cardiac function (Hulsmans et al., 2017; Nicolas-Avila et al., 2020). Whereas depletion of tissue macrophages leads to alteration in lipid storage (Cox et al., 2021; Gallerand et al., 2024) and adrenal macrophages have been implicated in metabolic homeostasis (Dolfi et al., 2022; Xu et al., 2024), the body composition of adult *Fireko* mice was not distinguished from WT (**Figure S2A**). Rather, resident macrophages appear essential to mitigate age-dependent neuropathology (Chadarevian et al., 2024; Kiani Shabestari et al., 2022; Munro et al., 2024) or tissue injury, for example in radiation-induced injury in the salivary gland or myocardial ischemia models (McKendrick et al., 2023; Weinberger et al., 2024) and the renal injury model in the current study.

### Repopulation of vacant territories in Fireko mice by wild-type bone marrow cells

The vacant macrophage territories of *Fireko* mice were occupied selectively by donor cells following intra-peritoneal WT bone marrow cell transfer at weaning. The expression of the FusionRed transgene in donor cells provides an example of the potential experimental application of this model to generate resident tissue macrophage-specific chimerism. Donor cells do not make any detectable contribution to bone marrow or circulating blood monocytes in this model. Repopulation of a vacant microglial niche has previously been demonstrated following neonatal adoptive transfer of monocytes purified from bone marrow (Bastos et al., 2025; Bennett et al., 2018). Whilst we cannot exclude a contribution from donor-derived monocytes trafficking from the peritoneum, at the least the tissue macrophage populations once established are maintained without monocyte recruitment. Based upon these observations and related findings in rats (Carter-Cusack et al., 2024; Keshvari et al., 2021; Sehgal et al., 2023) and chickens (Garceau et al., 2015) it seems likely that bone marrow contains a CSF1R-dependent macrophage progenitor population that gives rise directly to resident tissue macrophages. Future experiments will explore the relationship to the clonogenic monocyte/macrophage restricted progenitor defined by Hettinger *et al*. (Hettinger et al., 2013). Hematopoietic stem and progenitor cells and/or committed progeny have been shown to migrate in the circulation and lymphatics and to be capable of giving rise directly to tissue-resident myeloid cells (Massberg et al., 2007). One interesting question is whether the macrophage-depleted peritoneal cavity provides a unique trophic niche for expansion of these cells that might be exploited therapeutically. -

## Conclusion

Analysis of the *Fireko* mice has revealed surprising levels of redundancy in mononuclear phagocyte biology, in the function of the enhancer, the expression of CSF1R in progenitors and monocytes and in the roles of resident macrophages in tissue development and homeostasis. It is likely that tissue resident macrophages will prove to be more important to innate and acquired immune responses and repair following tissue injury.

## Materials and Methods

### Animals

The 418bp deletion containing the Fms intronic regulatory element (FIRE) to produce the *Csf1r*^ΔFIRE^ allele was generated using CRISPR-Cas9 in C57Bl/6J x CBA/J F1 zygotes (Rojo et al., 2019). The allele was backcrossed for 10 generations to C57BL/6J, then transferred to Australia, where it was crossed to *Csf1r*-EGFP reporter mice (Sasmono et al., 2003) on the C57Bl/6J/Arc background. Residual non-C57BL/6J contributions to the genome in this line were found to partly mitigate perinatal lethality and hydrocephalus (Preprint;(Taylor et al., 2025)). To restore resilience and generate larger cohorts for functional analysis, *Csf1r*^ΔFIRE/+^ mice were mated with wild-type CBA/J mice (Animal Resource Centre/WA) and F1 mice were crossed to generate homozygotes (*Fireko*) on the F2 background. The *Csf1r-*FusionRed knock-in reporter transgenic line was described elsewhere (Grabert et al., 2016) and is also maintained on the C57Bl/6J/Arc background.

We used CRISPR/Cas9 to mutate the AP1 binding site in the *Csf1r* gene. A sgRNA, aactgattcacagcctctga-ggg, directly targets the AP1 tgattca motif and a ssODN donor (tgccagcaatgtgtttccgcccacacaggccgggggcgcctgccaggccctcagaggctgTCTAGAgttctcacttccccccttcccccctatttcaagcctgggaaaaatgctgacaccacacag) with 60nt homology flanking a TCTAGA sequence to substitute this AP1 site was designed. In addition to destroying the AP1 site this integrates an XbaI restriction enzyme site used for rapid genotyping of founders.

The sgRNA was produced as an Alt-R crRNA (IDT) oligo and resuspended in sterile RNase free injection buffer (Tris HCl 1 mM, pH 7.5, EDTA 0.1 mM) and annealed with trans-activating crispr RNA (tracrRNA; IDT) by combining 2.5 μg crRNA with 5 μg tracrRNA and heating to 95 °C. The mix was left to slowly cool to room temperature (RT). After annealing the complex an equimolar amount was mixed with 1000 ng Cas9 recombinant protein (NEB; final concentration 20 ng/μL) and incubated at RT for 15 min, before adding Cas9 messenger RNA (mRNA) (final concentration; 20 ng/μL) and the ssODN polyacrylamide gel electrophoresis (PAGE) purified repair template (IDT; final concentration 50 ng/μL) in a total injection buffer volume of 50 μL. The injection mix was centrifuged for 10 min at RT and the top 40 μL removed for microinjection.

The mix was microinjected into one-day single cell mouse embryos (C57BL/6JOlaHsd). The zygotes were cultured overnight, and the resulting 2 cell embryos implanted into the oviduct of day 0.5 post-coitum pseudopregnant mice. Genotyping was performed using the Sigma redextract-n-amp tissue PCR kit. In brief, ear clips were digested then PCR performed at an annealing temperature of 62 °C with forward (5’ ctgtcactgtgtaggaagggt) and reverse primers (5’ gaaaccgttgtgtcatcccc). The DNA was then digested with 2 μL CutSmart Buffer (NEB, R0137L) and 1 μL XbaI enzyme (NEB) at 37 °C for 1 h. The products were analysed on a Qiaxcel automated gel system. A candidate founder was Sanger sequenced to confirm the desired genetic change, and a colony established and transferred to Australia.

All animal experiments in Australia were approved by The University of Queensland Health and Sciences Ethics Committee and performed in accordance with the Australian Code of Practice for the Care and Use of Animals for Scientific Purposes and Queensland Animal Care and Protection Act (2001). Animals were housed in individually ventilated cages with a 12 h light/dark cycle, and food and water available ad libitum.

### Cell isolation and culture

Peripheral blood was collected via cardiac puncture into EDTA-coated tubes and analysed using an automated hematology analyser, Mindray BC5000 Haematology Analyser (TRI Flow Cytometry Facility). Blood was then subjected to red blood cell lysis for 2 min in lysis buffer (150 mM NH_4_Cl, 10 mM KHCO_3_, 0.1 mM EDTA, pH 7.4), followed by two PBS washes and resuspension in FACS buffer (1x PBS with 2% FBS) for staining. Peritoneal cells were collected by lavage with 5 mL of PBS, centrifuged at 400 g for 5 min at 4°C, and resuspended in 1 mL of FACS buffer for staining. To generate elicited macrophages, a 4% sterile thioglycollate broth solution (Brewer) was prepared and administered as a single intraperitoneal (i.p.) injection on day 1. The control group received an equivalent volume of sterile saline via i.p. injection. Peritoneal cells were harvested by lavage on day 5.

Bone marrow (BM) cells were prepared by flushing femurs and tibias with 10 mL of PBS into a tube on ice, then centrifuging at 400 g for 5 min at 4°C. The pellet was resuspended in 1 mL of FACS buffer for staining. Following red cell lysis, single-cell suspensions were stained with fluorophore-conjugated antibodies on ice and analysed on an LSR Fortessa, (BD, Australia). Cell-surface markers used to define hematopoietic cell types and antibodies are provided in Supplemental Table 1. Data were analysed using FlowJo (BD). Cell counts were determined using a Mindray BC5000 Haematology Analyser (TRI Flow Cytometry Facility).

BM cells were seeded at 50,000 cells per well in the 96-well plate in 200 µL of complete medium (RPMI 1640 + 10% FBS, 25 U/mL penicillin, and 25 μg/mL streptomycin (Gibco, Thermo-Fisher, Australia) with added growth factors including CSF1 (100 ng/mL), CSF2 (100 ng/mL), FLT3L (20 ng/mL), KITL (20 ng/mL), IL3 (1:25 dilution), IL4 (20 ng/mL), IFN-γ (20 ng/mL), and CSF3 (100 ng/mL). Cells were then incubated at 37°C in a humidified atmosphere with 5% CO₂. On day 7, resazurin (10ul, 25ug/ml) was added to each well. Following incubation for 1–4 hours at 37°C, fluorescence was measured using an Omega Pherastar plate reader (BMG Labtech) with excitation and emission wavelengths set to 530–540 nm and 585–595 nm, respectively. Gain adjustments (1500–1800) were applied to optimise signal detection. The fluorescence signal, which is proportional to cell number, was used to evaluate cell viability.

To assess CSF1R expression, BM cells grown in different growth factors were washed and detached by adding 800 µL of cold PBA buffer (1x PBS containing 1% BSA and 0.05% sodium azide) for 3 minutes. Cells were collected in FACS tubes and centrifuged at 400 × g for 5 minutes. The supernatant was aspirated, leaving ∼25 µL, to which 25 µL of antibody cocktail containing PE-Cy7-labeled anti-CD115 and other relevant antibodies was added. Cells were vortexed, incubated on ice for 30 minutes on a shaking platform, and washed twice with 800 µL of PBA buffer. Following staining, cells were resuspended in 100 µL of PBA buffer and viability was assessed using 7-AAD staining. The samples were incubated with 10 µL of 7AAD for 5–10 minutes before analysis. Flow cytometry was performed using standard excitation/emission settings appropriate for the fluorophores, and CSF1R expression (CD115) was quantified as part of a multiparametric panel including CD45, CD11b, and F4/80.

For bulk cultures to generate bone marrow-derived macrophages (BMM), ca. 10^7^ BM cells were seeded on 100 mm square bacteriological plastic (Sterilin, Thermo-Fisher, Australia) in 25 ml of complete medium (RPMI + 10% FBS, 25 U/mL penicillin, and 25 μg/mL streptomycin (Gibco, Thermo-Fisher, Australia) and differentiated for 7 days with the addition of recombinant CSF1-Fc (100 ng/ml) or recombinant mouse CSF2 (GM-CSF, R&D Systems, Minneapolis, Mn, USA; 50 ng/ml). For osteoclast culture bone marrow cells were harvested as described above and seeded at 5 x 10^4^ cells/ml in 12 well tissue culture plates in complete medium with CSF1 (100 ng/ml) for 3 days. From day 3 the medium was replaced daily with medium containing CSF1-Fc (100 ng/ml) and RANKL (R&D Systems, 40 ng/ml) until day 7. Tartrate-resistant acid phosphatase (TRAP) staining was performed in situ using a kit (Cat #387A, Sigma, Australia) according to the manufacturer’s instructions.

### Whole mount imaging

Whole mount imaging of the *Csf1r*-EGFP and *Csf1r-*FusionRed reporters was performed using an FV3000 microscope (Olympus) and FLUOVIEW (Olympus) software. Tissue was harvested and kept in ice-cold PBS until imaging. Tissue was placed flat on a glass petri dish. Z-stack images were taken using the AF488 and AF594 lasers and combined to produce maximum intensity projections. Images were taken at 10x or 20x magnification and processed on ImageJ software v1.54.

### Immunofluorescence staining

For cryosections, spleens were dissected and immersion fixed in 4% PFA for 4h, before cryoprotection in 15% sucrose in PBS overnight and then 30% sucrose in PBS for 48 h. Spleens were then embedded in OCT, frozen and sectioned at 10μm using a ThermoFisher HM525NX cryostat. For staining, sections were rehydrated in Tris-buffered saline (TBS; 10mM Tris, 150mM NaCl, pH8.0) for 10 min and washed 3 x min in TBST (0.01% Tween-20 in TBS). Blocking buffer (3% BSA, 5% serum in TBST) was applied for 30 min at RT, before sections were incubated with primary antibodies: rat anti-CD169 (1:100, Biolegend, 142402) and rabbit anti-CD209B (1:150, Abcam, ab308457), diluted in TBST for 1.5h at RT. Sections were washed 3 x 5 min TBST before incubation in secondary antibodies: donkey anti-rabbit AF647 (1:500, Invitrogen, A31573) and goat anti-rat AF594 (1:400, Invitrogen, A11007) diluted in TBST for 45 min protected from light. Sections were washed 3 x 5 min in PBS, stained with DAPI and mounted for imaging.

### F4/80 immunohistochemistry

10 µm paraffin sections were dewaxed, rehydrated and incubated for 30 mins in 10 mM Tris/EDTA (pH 9) buffer maintained at 90-100°C using a microwave for antigen retrieval. Slides were cooled on ice for 10 min, incubated in 0.3% hydrogen peroxide (Sigma) for 15 min to block endogenous peroxidases and washed in TBS. Slides were incubated in 1% BSA in TBS for 30 min followed by incubation with primary antibody (anti-F4/80, 1:1000, rabbit, Abcam) for 1.5 hr in a humidified chamber. Slides were washed with TBS, incubated sequentially with HRP secondary reagent (anti-rabbit, Agilent) for 30 min, DAB Peroxidase substrate (Agilent) for 5 mins and haematoxylin for 40 sec then dehydrated and mounted. Imaging was performed on an Olympus BX50 microscope, using brightfield settings.

### Quantification of mRNA expression

Tissues for qRT-PCR analysis were snap frozen on dry ice and stored in a -80°C freezer. Approximately 50–100 mg of frozen tissue was homogenized in 1 mL of cold TRIzol™ reagent using a FastPrep homogenizer (40 seconds at 5 m/s). The homogenates were centrifuged at 12,000 × g for 5 minutes at 4°C, and the clear supernatant was transferred to a fresh tube.

Following a 5-minute incubation to dissociate nucleoprotein complexes, 0.2 mL of ice-cold chloroform was added per mL of TRIzol™ reagent, and the mixture was inverted 5–10 times, incubated for 3 minutes, and centrifuged at 12,000 × g for 15 minutes at 4°C. The RNA-containing aqueous phase was carefully transferred to a new tube, mixed with 0.5 mL of isopropanol, and incubated for 10 minutes to precipitate RNA. The RNA pellet was washed twice with 75% ethanol, air-dried for 5–10 minutes, and dissolved in 20–50 µL of RNase-free water. Dissolved RNA was incubated at 55–60°C for 10–15 minutes to ensure complete solubilization. RNA quantity was measured using a NanoDrop Spectrophotometer (Thermofischer Scientific).

Complementary DNA (cDNA) was synthesised from 1 µg of total RNA using the SensiFAST cDNA Synthesis Kit (Bioline). RNA was first treated with DNase I (Sigma) to remove genomic DNA contamination by incubating the RNA with DNase I (0.2 µL) and 10× buffer (1 µL) in a total volume of 10 µL at room temperature for 15 minutes, followed by inactivation with 1 µL of 25 mM EDTA at 65°C for 10 minutes. Reverse transcription was performed in a 20 µL reaction containing 11 µL RNA, 4 µL 5× TransAmp buffer, 0.25 µL reverse transcriptase, and 4.75 µL RNase-free water. Reactions were incubated at 24°C for 10 minutes (primer annealing), 42°C for 15 minutes (reverse transcription), and 85°C for 5 minutes (enzyme inactivation). The resulting cDNA was diluted 1:10 with RNase-free water for subsequent quantitative PCR analyses and stored at -20°C.

Gene expression analysis was performed using quantitative real-time PCR (qRT-PCR) with SYBR Green PCR Master Mix. Each reaction was conducted in a 10 µL volume containing 2 µL of diluted cDNA, 2 µL of forward and reverse primer mix (1–2 µM final concentration), 5 µL of SYBR Green Master Mix, and 1 µL of RNase-free water. Thermal cycling was carried out on a real-time PCR system with the following conditions: initial denaturation at 95°C for 2 minutes, followed by 40 cycles of 95°C for 15 seconds, 60°C for 1 minute, and a melt curve analysis to confirm specificity. Relative expression levels of target genes were normalised to the expression of a housekeeping gene using the ΔΔCt method. All reactions were performed in triplicate to ensure reproducibility.

Primer sequences used were:

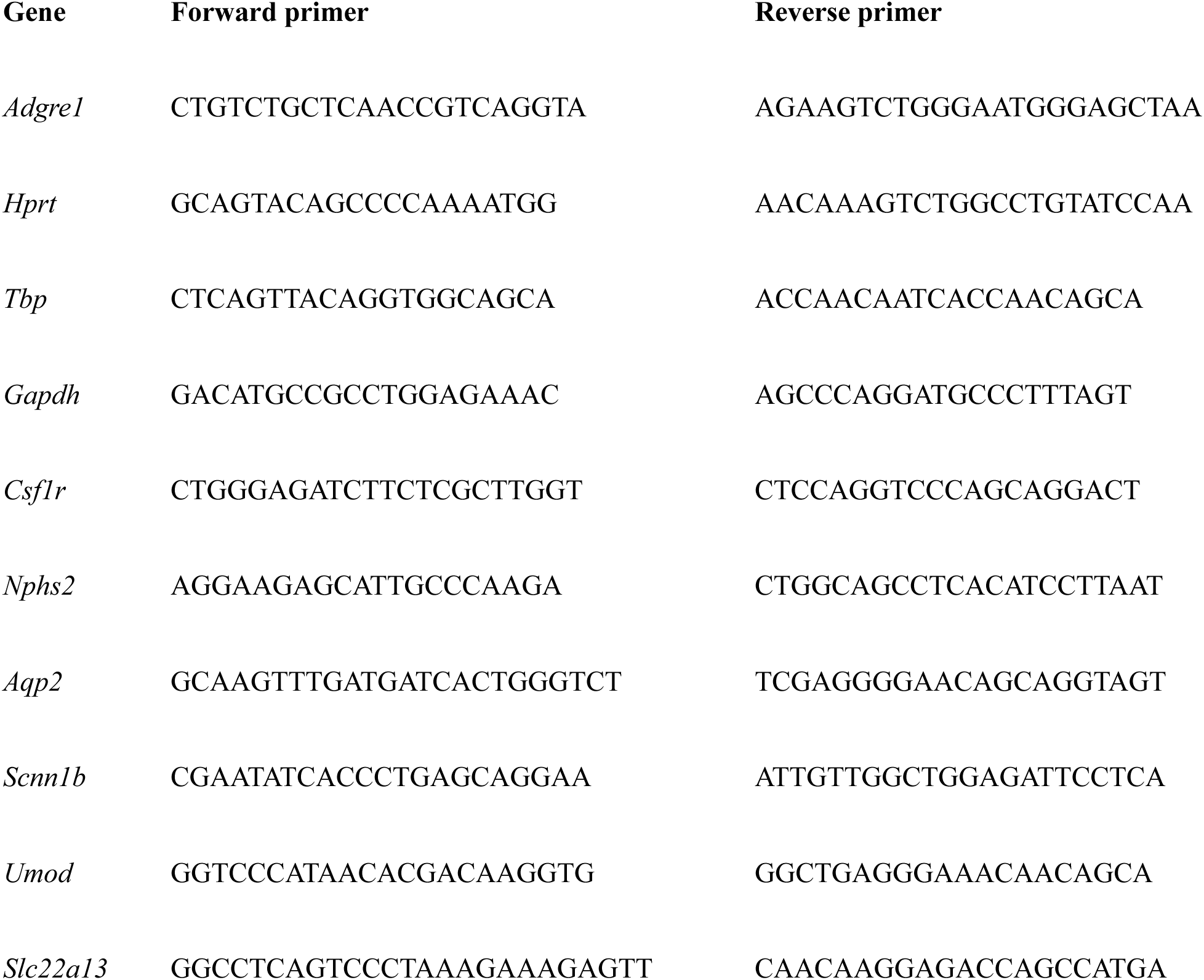

### Treatment with CSF1 in vivo

To assess *in vivo* response to CSF1, mice were injected with a recombinant human-CSF1 mouse Fc conjugate (Novartis, Switzerland). Mice received one injection per day of 1 mg/kg of CSF1-Fc for 4 days between Zeitgeber time (ZT) 2-3. Mice were placed in the Minispec LF5OH (Brucker) at ZT2 to measure their body composition and blood glucose prior to the first injection. Their body weight was monitored daily. On day 5, body composition and blood glucose were again measured and mice were euthanised with CO_2_ prior to tissue collection.

### Thioglycollate-elicited peritoneal macrophages

1ml of thioglycolate broth was injected into the peritoneal cavity of adult male WT and *Fireko* mice. Peritoneal cells were harvested by lavage after 5 days.

### Adoptive transfer of wild-type bone marrow cells

Bone marrow cells were harvested from the tibia and femur of *Csf1r*-FusionRed female mice. The bones were flushed with a solution containing 0.9% sodium chloride and 2% heat-inactivated FBS. The cell suspension was then filtered through a 70µm cell strainer and centrifuged at 500g for 5 mins at 5°C. The pelleted cells were resuspended in 1x sterile PBS supplemented with 2% heat-inactivated FBS (10mL per bone) for cell counting using a haemocytometer (Reichert, U.S.A). Following counting, cells were centrifuged and resuspended in 1x sterile Dulbecco’s phosphate-buffered saline (DPBS, Gibco, 10010-023). 2 × 10^7^ *Csf1r*-FusionRed cells were injected i.p. at weaning administered intraperitoneally (i.p.) and recipients were analysed 12 weeks post transfer.

### Adenine chronic kidney disease model

Male and female 8 week old wild-type and *Fireko* mice were exposed to 0.2% adenine in the drinking water for 4 weeks as described (Diwan et al., 2018). Bodyweights were measured weekly. At sacrifice blood was taken for measurement of urea, creatinine and phosphate by University of Queensland Veterinary Pathology and tissues processed for histology and immunohistochemistry.

Kidneys were fixed in 4% PFA for 4 hours, dehydrated and embedded in paraffin wax. Thin sections (5 μm) of the kidney were cut and stained with H&E. An Olympus VS120 Slidescanner Microscope (TRI) for imaging was used, and image morphometry was analysed by a veterinary pathologist (Dr Rachel Allavena) who was blinded to sample identity. Percent tubulointerstitial atrophy and percent tubules with casts/cell debris were assessed morphometrically in four fields/kidney at ×20 digitally scanned magnification. Apoptosis was counted using structural characteristics in the tubular epithelium of four fields/kidney at ×20 digitally scanned magnification. The inflammatory infiltrate was assessed on a 4 point scale: 1 = normal; 2 = little infiltrate (<10% of field), 3 = moderate infiltrate (10–50% of field), and 4 = extensive infiltrate (>50% of field, and usually >>50%/field). Glomerular pathology (sclerosis, thickening of Bowman’s capsule) was assessed as the mean of individual glomeruli from four scanned cortical fields per kidney section. Loss of brush borders was assessed using periodic acid-Schiff (PAS). Paraffin-embedded tissue sections were deparaffinized and hydrated to deionized water. Sections were incubated in periodic acid solution (1 g/dL) (Sigma Aldrich, 3951) for 5 minutes at room temperature and rinsed in several changes of distilled water. Slides were then immersed in Schiff’s reagent (Sigma Aldrich, 3952) for 15 minutes at room temperature, followed by washing in running tap water for 5 minutes. Collagen deposition was assessed by incubating sections with a Picro-sirius Red solution (0.2 g Sirius Red F3B in 500 mL saturated picric acid) for 1 hour in a glass or plastic container. After staining, sections were washed in two changes of acidified water (5 mL glacial acetic acid in 1 L distilled water) with gentle agitation (20–22 seconds per wash). Sections were further washed in 100% ethanol for 5 minutes twice to prevent cytoplasmic destaining. After dehydration through three changes of 100% ethanol, sections were cleared in xylene and mounted using DPX Mountant (Sigma Aldrich, 06522). Under bright-field microscopy, collagen appeared red on a pale yellow background, with birefringent collagen fibers visible under polarised light. Slides were then scanned using the Olympus VS120 Slidescanner Microscope (TRI).

### Cardiac ultrasound

Cardiac ultrasound was performed using the Vevo 2100 Imaging System. Mice were anesthetised and then positioned on a platform with electrocardiogram and temperature monitoring and prepared through hair removal for optimal imaging. Probe selection was tailored to the size of the animal and its heart (e.g., MS400 for mice), ensuring optimal image acquisition. The parasternal long and short axis views in echocardiography were utilised to visualise cardiac anatomy and assess function. The parasternal long axis (PLAX) view provided a sagittal section of the heart, revealing structures such as the left and right ventricles, left atrium, aorta, mitral and aortic valves, and pulmonary artery. This view was employed to measure parameters including ejection fraction and cardiac output. To obtain this view, the probe was positioned vertically and rotated slightly counterclockwise. In contrast, the parasternal short axis (PSAX) view offered a transverse section of the heart, enabling visualisation of the ventricles, intraventricular septum, papillary muscles, and other structures. This view was achieved by rotating the probe 90° clockwise from the PLAX orientation. These complementary views formed the basis of a comprehensive evaluation of cardiac function, including wall motion, valve function, and blood flow dynamics. The Simpson method was employed to assess left ventricular (LV) volumes and stroke volume by measuring the LV at multiple planes during diastole and systole. Short-axis images were acquired at three levels: near the apex (SimpAreaDist), at the mid-section near the papillary muscle (SimpAreaMid), and just below the base of the aorta (SimpAreaProx). At each level, the endocardial border was traced in both diastole and systole. Additionally, a parasternal long-axis view was obtained to measure the LV length from the aortic annulus to the endocardial border at the apex during diastole and systole. These measurements were used to calculate diastolic and systolic volumes, as well as the stroke volume, using the Vevo 2100 Imaging System’s calculation tools.

## Acknowledgements

The generation of the mice used in this study was funded originally by a grant from the Medical Research Council (MRC) UK grant MR/M019969/1 to DAH. This work was supported by core support and direct funding from The Mater Foundation and NHMRC Investigator Grant #2007950 to DAH. We acknowledge input and expertise from the Biological Resources facility and the Preclinical Imaging, Microscopy, Histology and Flow Cytometry facilities of the Translational Research Institute (TRI). TRI is supported by the Australian Government.

