## Supplementary material for "Depletion and replacement of tissue resident macrophages in mice with germ-line deletion of a conserved enhancer in the *Csf1r* locus": Supplemtary Figures S1-S10

### Figure S1

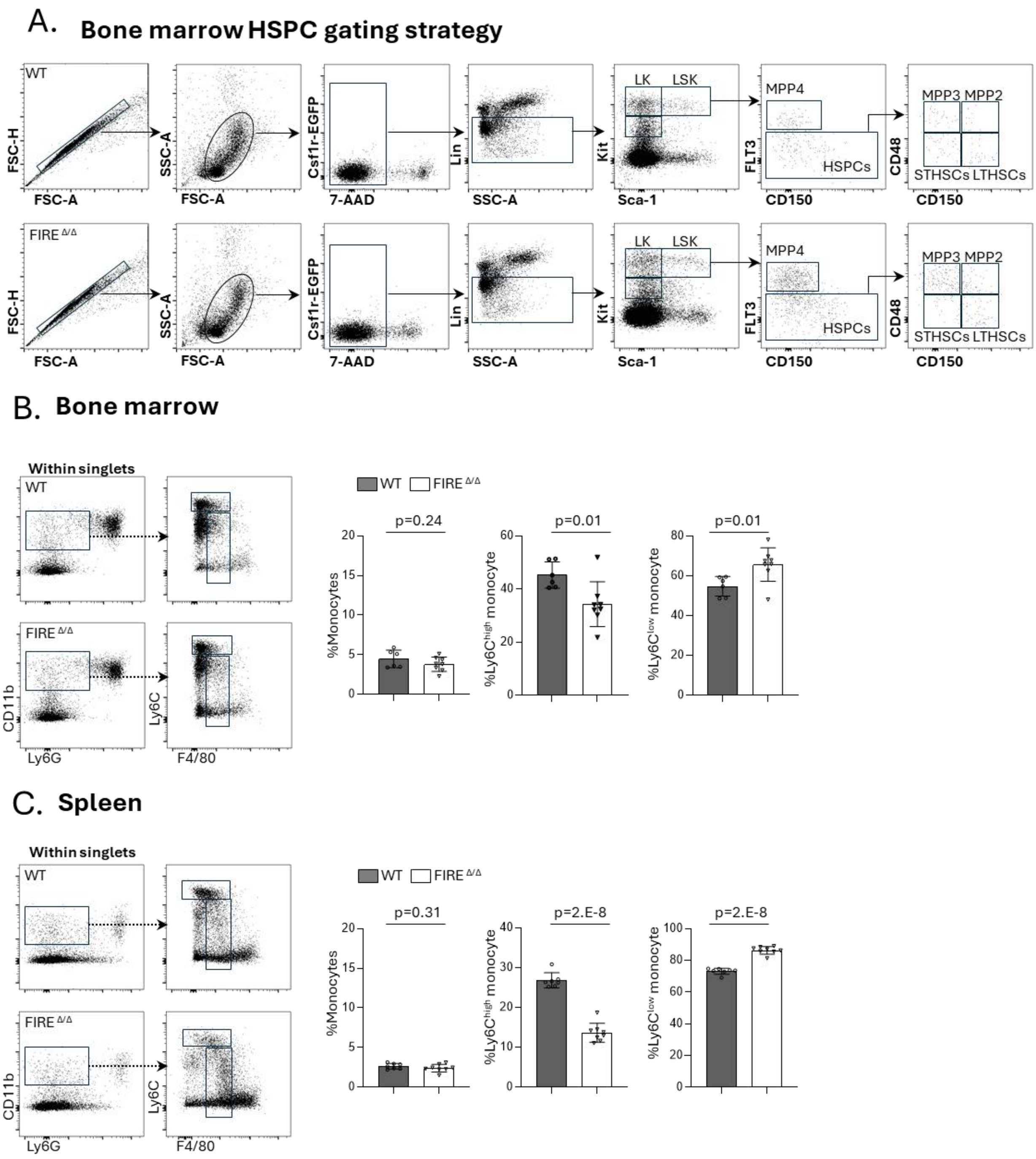


**Figure S1. Analysis of bone marrow progenitors and monocytes in WT and *Fireko* mice.**

(A) Gating strategy for quantitation of hematopoietic stem and progenitor cell populations shown in Figure 1. Panels show representative profiles for WT and *Fireko* mice. (B,C) Quantification of monocyte subpopulations in bone marrow and spleen. Monocytes are defined as Cd11b$^{+}$, Ly6G$^{-}$, F4/80$^{low}$ and further subdivide into Ly6C$^{high}$ and Ly6C$^{low}$

subpopulations as shown in typical flow cytometry profiles. Data show means and standard deviation. *p* values were determined by unpaired two-tailed t-test.

Figure S2

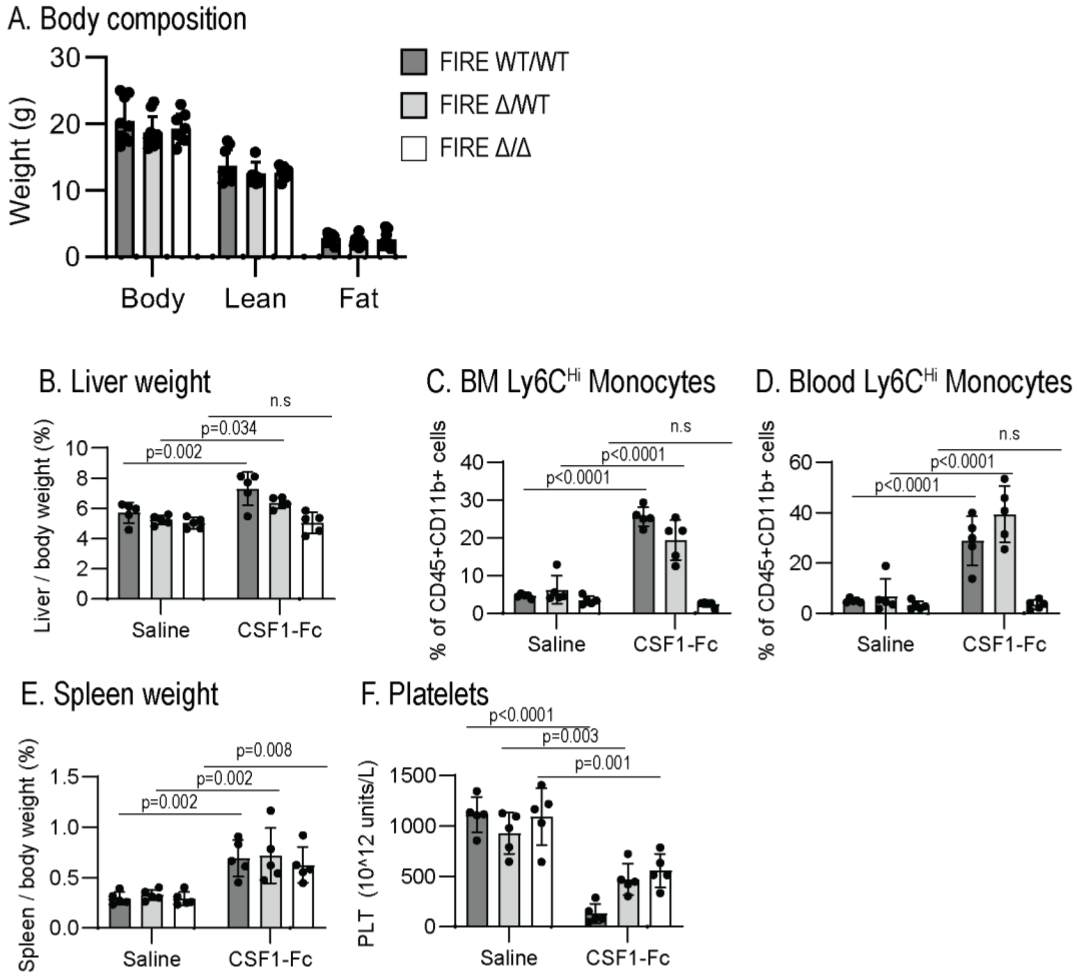


**Figure S2. CSF1 response in *Fireko* mice *in vivo.***

WT, *Csf1r*$^{\Delta FIRE/+}$ and *Fireko* mice on the C57BL/6J/CBA F2 background were injected with CSF1-Fc (1 mg/kg) or saline on each of 4 successive days and euthanised on day 5. (A) Prior to treatment, body composition was analysed using a Bruker Minispec. Each point is an individual animal. (B-F) C57/CBA mice were injected with CSF1-Fc (1 mg/kg) or saline on each of 4 successive days and euthanised on day 5. n = 5 per genotype (B) Liver/body weight ratio. Bone marrow (C) and peripheral blood (D) Ly6C$^{high}$ monocytes were gated as CD45$^{+}$/Ly6G$^{-}$/CD11b$^{+}$/Ly6C$^{high}$. (E) Spleen/body weight ratio.(F) Platelet counts were determined using Mindray analyser. Data are mean ± SD; two-way ANOVA with Sidak's multiple comparison test for Saline vs CSF1-Fc, n.s. not significant.

Figure S3

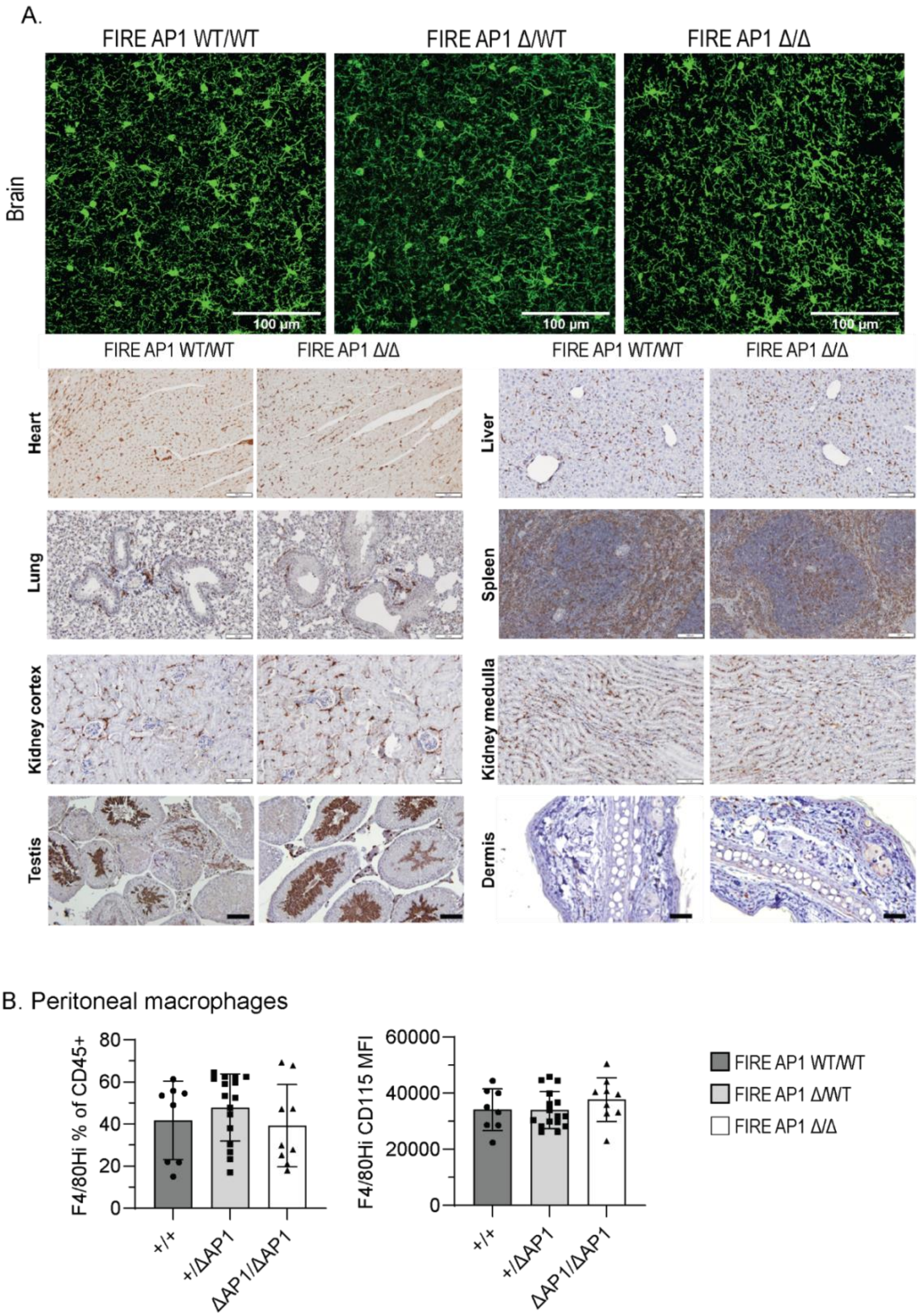


**Figure S3. *Csf1r* FIRE-AP1 mutation does not impact brain or peripheral macrophage populations.**

(A) Brain cortex, heart, liver, lung, spleen, kidney, testis and dermis sections from 9 week old *Csf1r*$^{+/+}$, *Csf1r*$^{+/\Delta AP1}$ or *Csf1r*$^{\Delta AP1/\Delta AP1}$ mice were stained for IBA1 (representative images). Scale bar in peripheral tissues = 50 µm. (B) Peritoneal cells from 3 week old *Csf1r*$^{+/+}$, *Csf1r*$^{+/\Delta AP1}$ or *Csf1r*$^{\Delta AP1/\Delta AP1}$ mice were analysed by flow cytometry. Resident peritoneal macrophages were gated as CD45$^{+}$/Cd11b$^{high}$/F4/80$^{high}$. MFI: median fluorescence intensity.

Figure S4

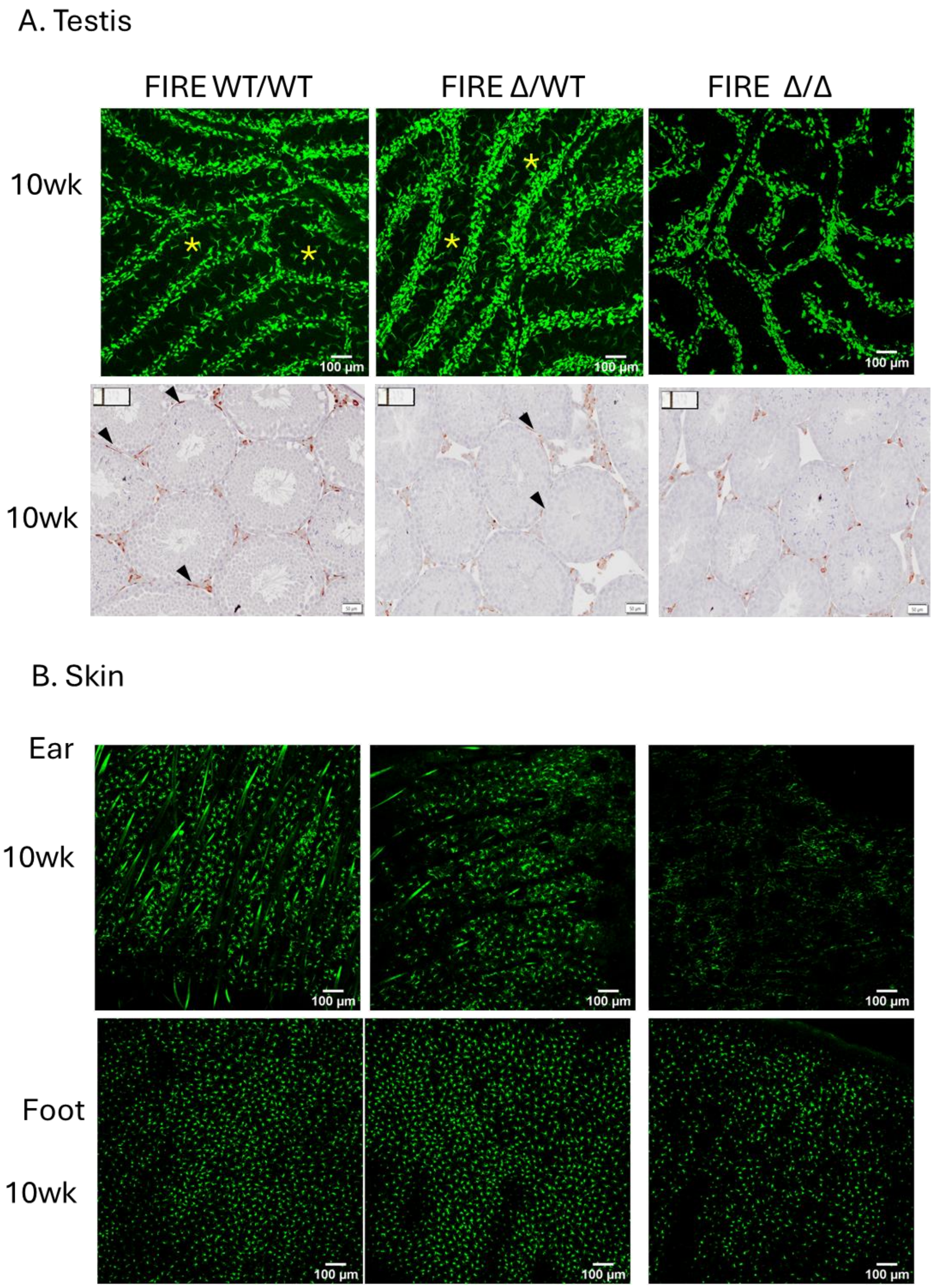
A. Testis
FIRE WT/WT
FIRE Δ/WT
FIRE Δ/Δ
10wk
100 µm
10wk
B. Skin
Ear
10wk
100 µm
Foot
10wk
100 µm

Figure S4

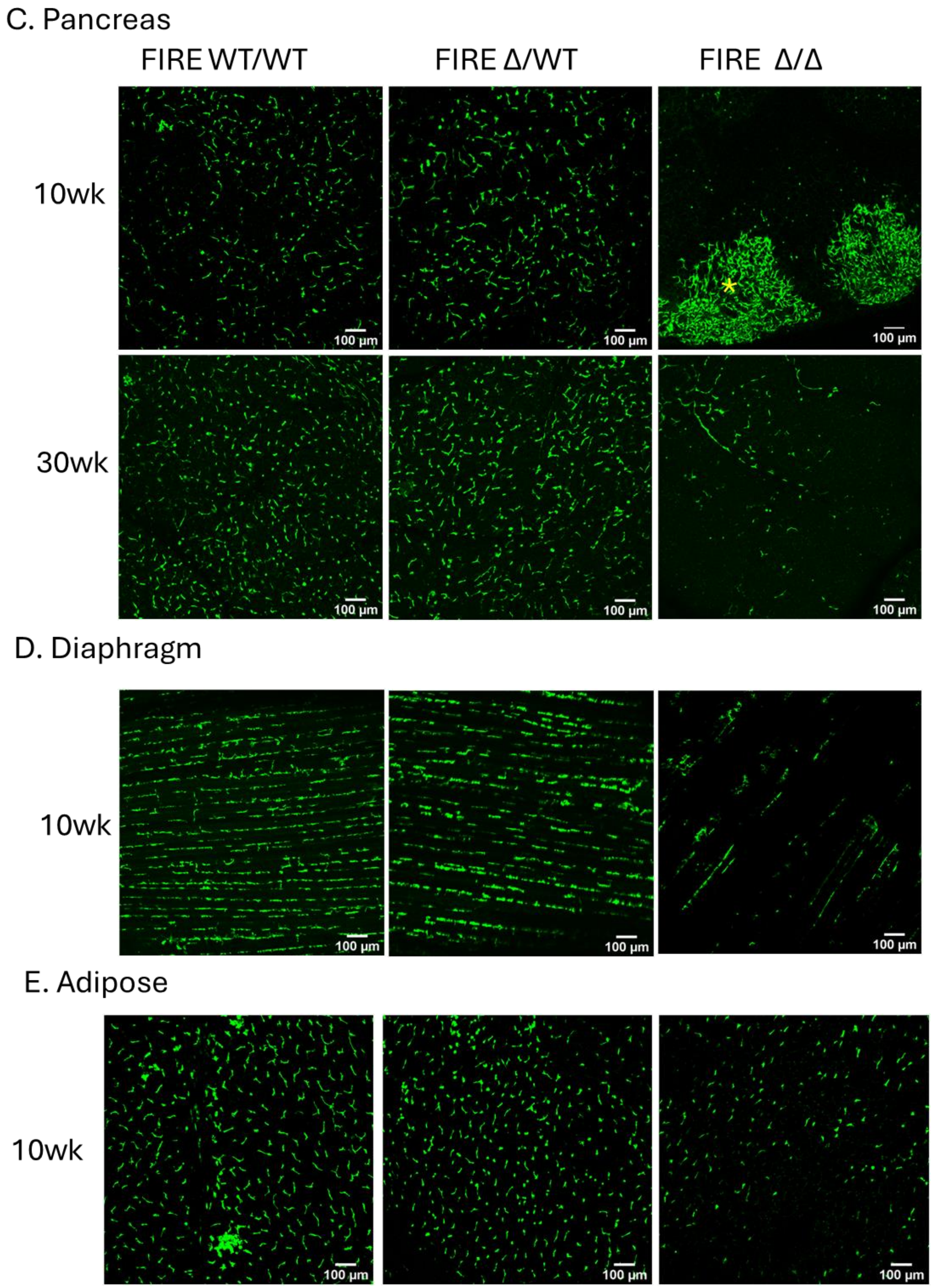
C. Pancreas
FIRE WT/WT
FIRE Δ/WT
FIRE Δ/Δ
10wk
30wk
100 µm
D. Diaphragm
10wk
100 µm
E. Adipose
10wk
100 µm

**Figure S4. Detection of tissue resident macrophages in *Fireko* mice.**

Representative whole mount images of *Csf1r*-EGFP in indicated tissues from adult male WT, *Csf1r*$^{\Delta FIRE/+}$ and *Fireko* mice. In the testis, note the GFP$^{+}$ macrophages spread on the surface of the tubules (asterisks) in WT and D/+ that are not detected in *Fireko* (D/D) whereas the interstitial macrophages associated with the interstitium are retained. The lower panel shows localisation of F4/80. The image of the *Fireko* pancreas at 10 weeks includes two lymphoid aggregates (asterisk) where EGFP$^{+}$ cells are retained, whereas no signal is detected in the acinar tissue.

Figure S5

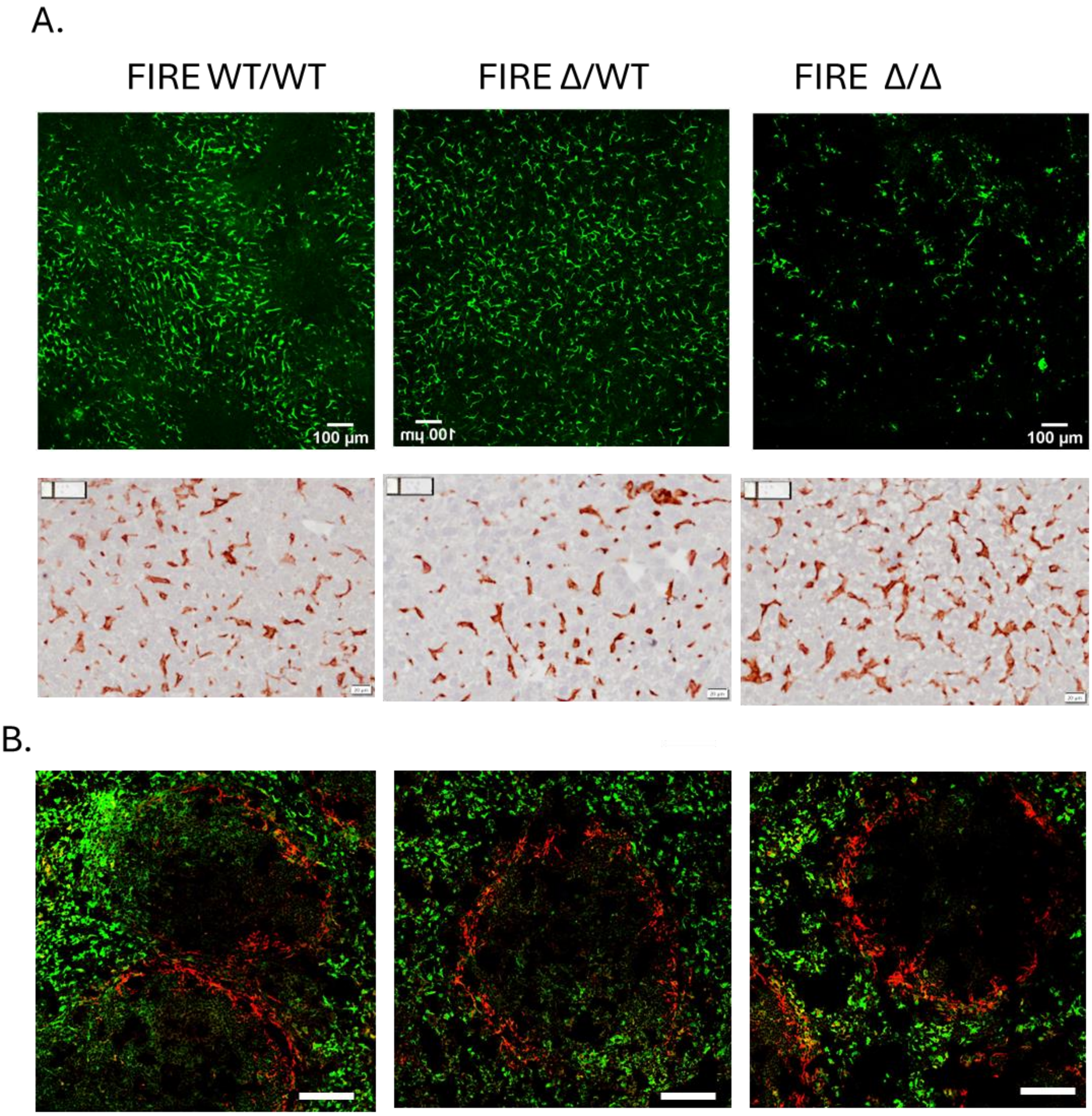


**Figure S5. Detection of tissue resident macrophages in *Fireko* mice.**

(A) Representative whole mount images of *Csf1r*-EGFP and F4/80 staining of liver from adult male WT, *Csf1r*$^{\Delta FIRE/+}$ and *Fireko* mice. Note the depletion of EGFP$^{+}$ capsular macrophages in *Fireko* (D/D) liver whereas F4/80$^{+}$ Kupffer cells are unaffected. B) Representative images of *Csf1r*-EGFP and CD169 (red) immunofluorescence localisation in spleen from adult male WT, *Csf1r*$^{\Delta FIRE/+}$ and *Fireko* mice. (C) F4/80 staining in adrenal gland sections from adult WT, *Csf1r*$^{\Delta FIRE/+}$ and *Fireko* mice.

Figure S6

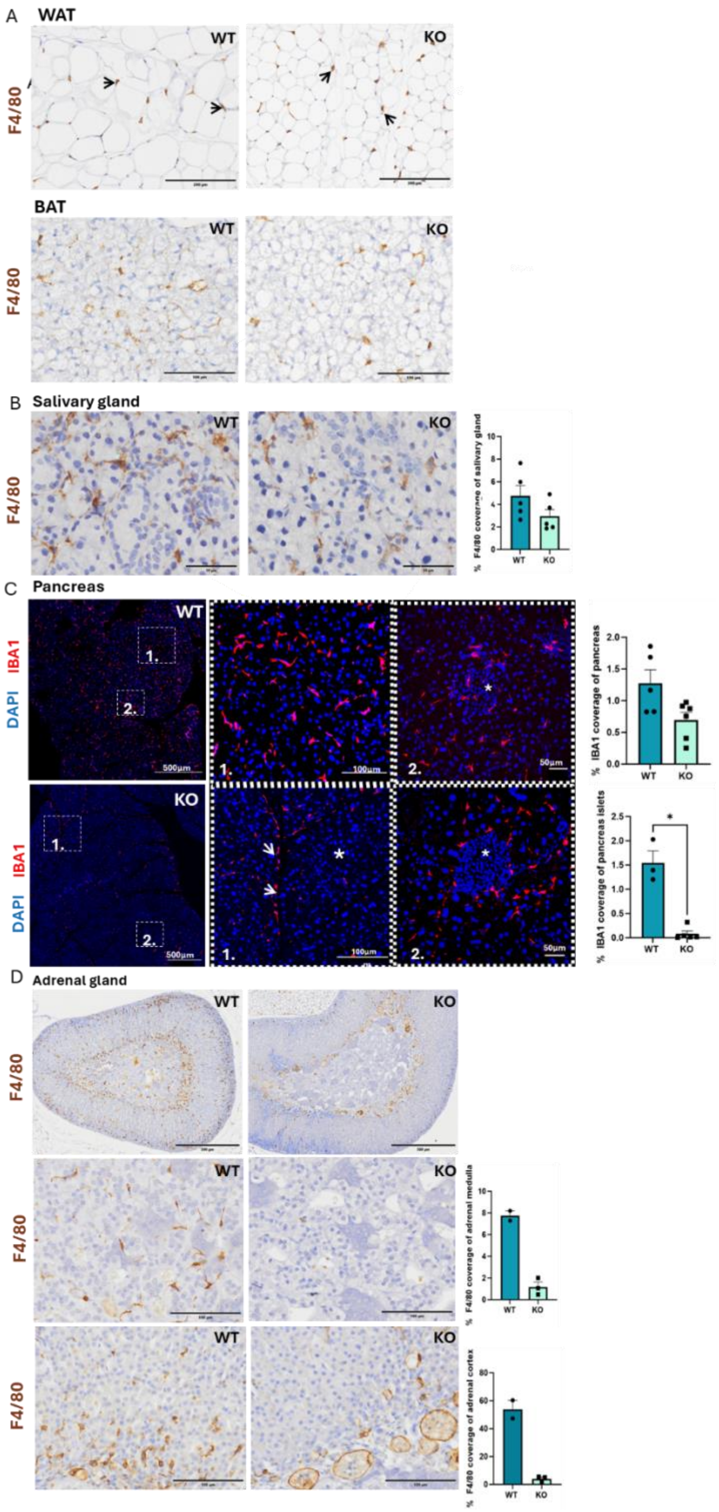
A
WAT
F4/80
WT
KO
BAT
B
Salivary gland
% F4/80 coverage of salivary gland
C
Pancreas
IBA1
DAPI
500µm
100µm
50µm
% IBA1 coverage of pancreas
% IBA1 coverage of pancreas islets
D
Adrenal gland
% F4/80 coverage of adrenal medulla
% F4/80 coverage of adrenal cortex

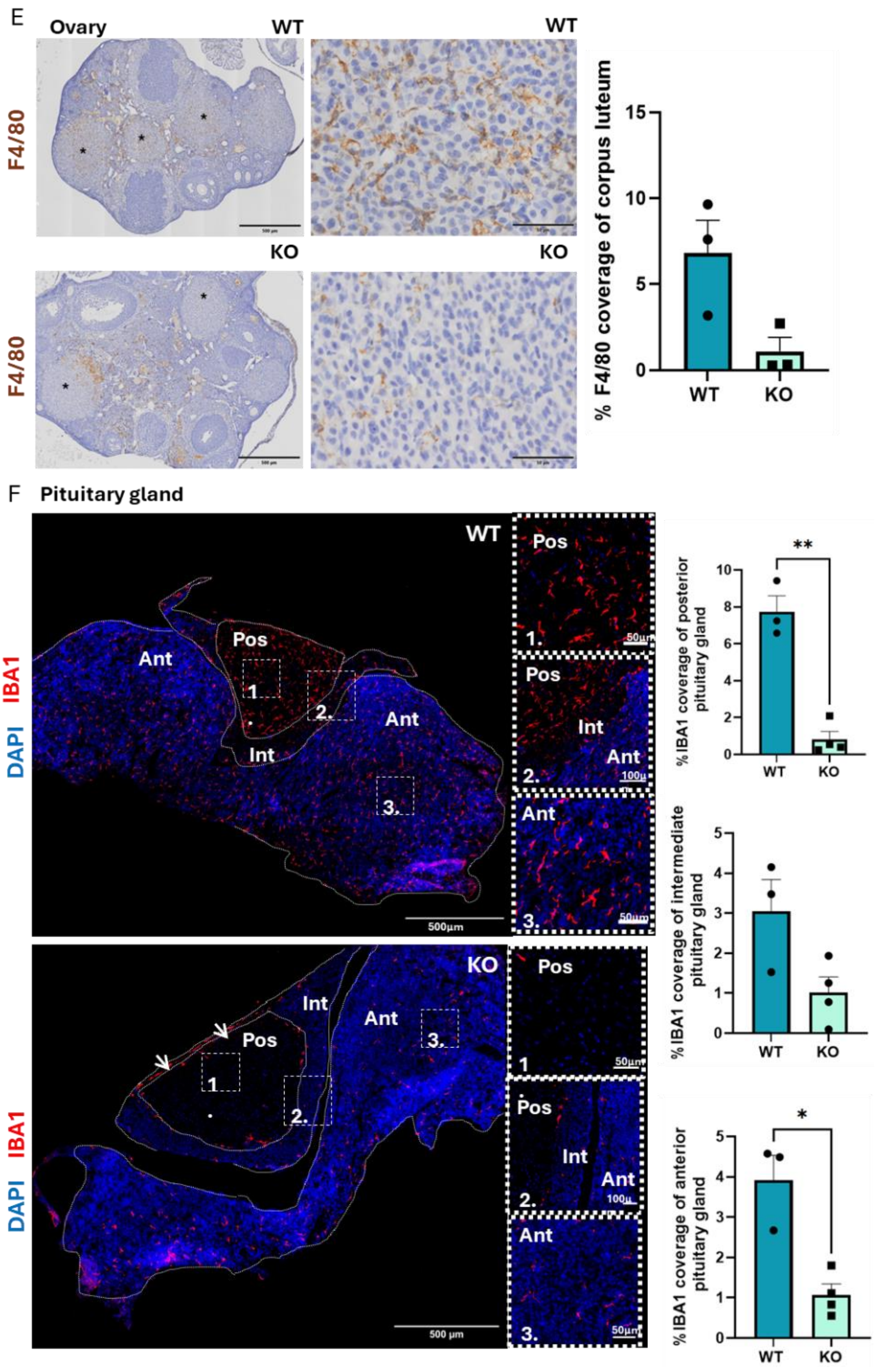


**Figure S6. Detection of tissue resident macrophages in *Fireko* mice.**

(A) Representative images of F4/80 staining of white adipose tissue and brown adipose tissue from adult WT and *Fireko* mice. Arrows highlight individual F4/80$^{+}$ interstitial macrophages. No quantitative diferennce was detected based upon genotype.

(B) Representative images of F4/80 staining of salivary gland from adult WT and *Fireko* mice and quantitation.  N=5 per group.

(C) Localisation of IBA1$^+$ macrophages in the pancreas of WT and *Fireko* mice. Panels show typical higher magnification images of exocrine and endocrine (islet) tissue (asterisks). % stained area was quantified; n=5 per group. Note that although there was not a significant difference between WT and *Fireko* in exocrine tissue, in the *Fireko* pancreas IBA1$^+$ cells appeared restricted to interlobular connective tissue (arrows).

(D) Representative images of F4/80 staining of adrenal gland tissue from adult WT and *Fireko* mice. Higher magnifications show the medulla and cortex. % stained area was quantified; n=3 per group.

(E). Representative images of F4/80 staining of ovaries from adult WT and *Fireko* mice. Higher magnifications show corpus luteum (asterisks). % stained area in corpora lutea was quantified; n=3 per group.

(F) Localisation of IBA1$^+$ macrophages in the pituitary of WT and *Fireko* mice. Low and high magnification images show the separate posterior, intermediate and anterior pituitary lobes as indicated. % stained area in each area was quantified; Each point is an individual mouse. Note that posterior pituitary parenchyma lacks detectable IBA1$^+$ cells; IBA1$^+$ cells are retained only on the surface (arrows).

### Figure S7

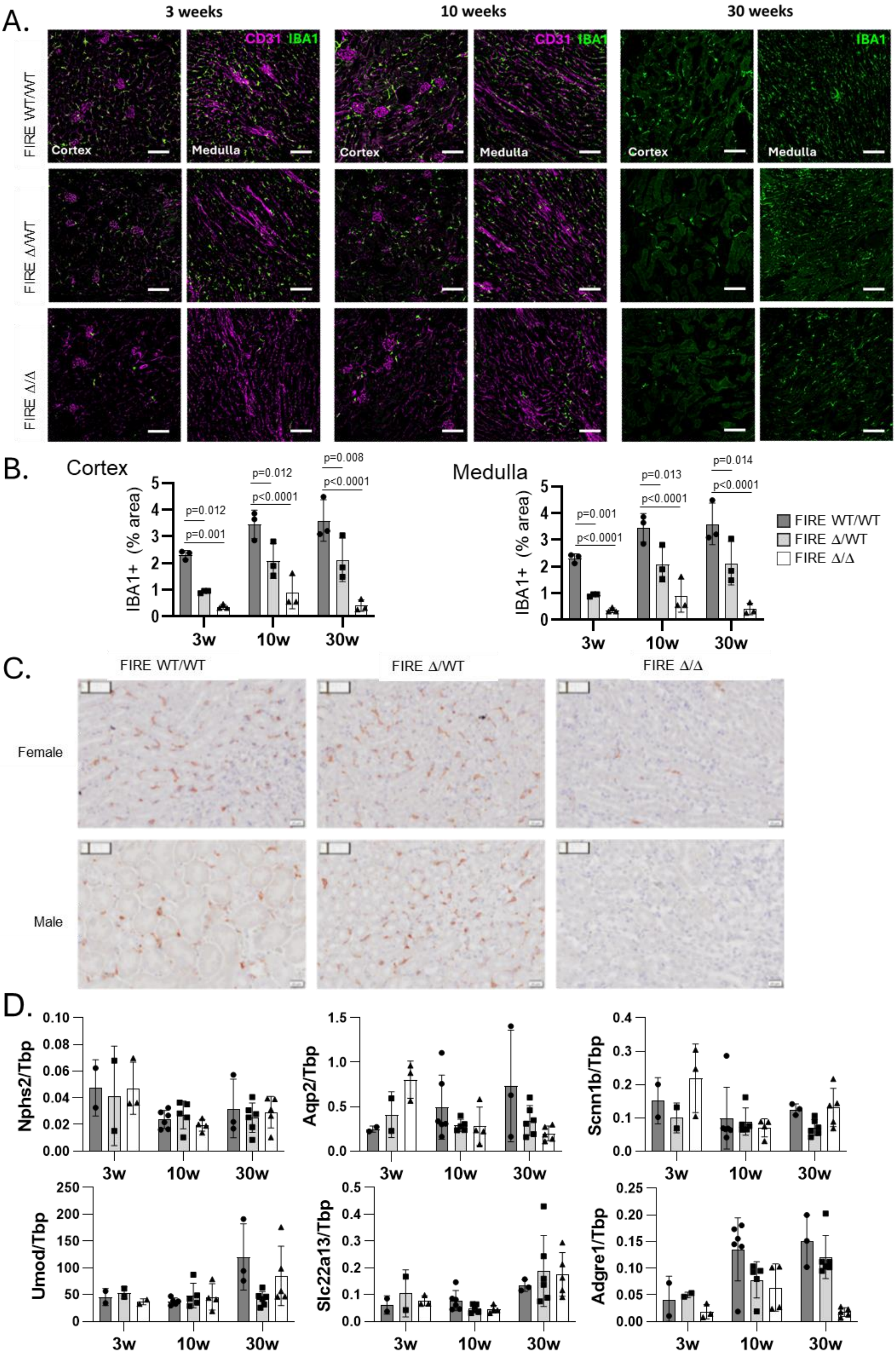

A.
3 weeks
10 weeks
30 weeks
CD31 IBA1
IBA1
FIRE WT/WT
FIRE Δ/WT
FIRE Δ/Δ
Cortex
Medulla
B.
Cortex
Medulla
IBA1+ (% area)
p=0.012
p=0.001
p=0.012
p<0.0001
p=0.008
p<0.0001
p=0.001
p<0.0001
p=0.013
p<0.0001
p=0.014
p<0.0001
3w
10w
30w
FIRE WT/WT
FIRE Δ/WT
FIRE Δ/Δ
C.
Female
Male
D.
Nphs2/Tbp
Aqp2/Tbp
Scnn1b/Tbp
Umod/Tbp
Slc22a13/Tbp
Adgre1/Tbp

**Figure S7. Renal macrophage deficiency in *Fireko* mice**.

(A) Representative images of sections from WT, *Csf1r*$^{\Delta FIRE/+}$ and *Fireko* mice at 3, 10 or 30 wks of age stained for IBA1 and CD31. Scale bar 100μm.

(B) IBA1$^{+}$ area in the cortex and medulla was quantified using image analysis. Data show mean and standard deviation, each datapoint is an individual mouse. 2 way Anova with Tukey's multiple comparison test.

(C) Representative images of kidney sections from WT, *Csf1r*$^{\Delta FIRE/+}$ and *Fireko* mice at 10 wks of age were stained for F4/80.

(D) Kidney mRNA expression of *Nphs2, Aqp2, Scnn1b, Umod, Slc22a13* and *Adgre1* was determined by qRT-PCR at 3, 10 and 30 wks of age as indicated

Figure S8

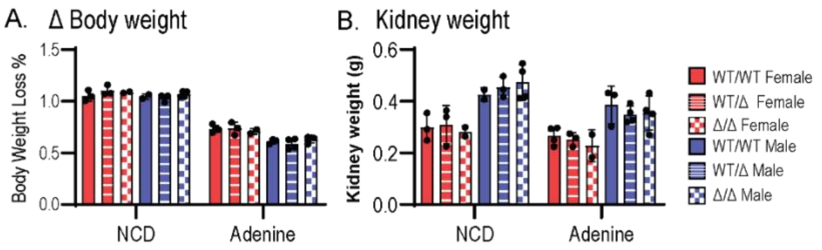


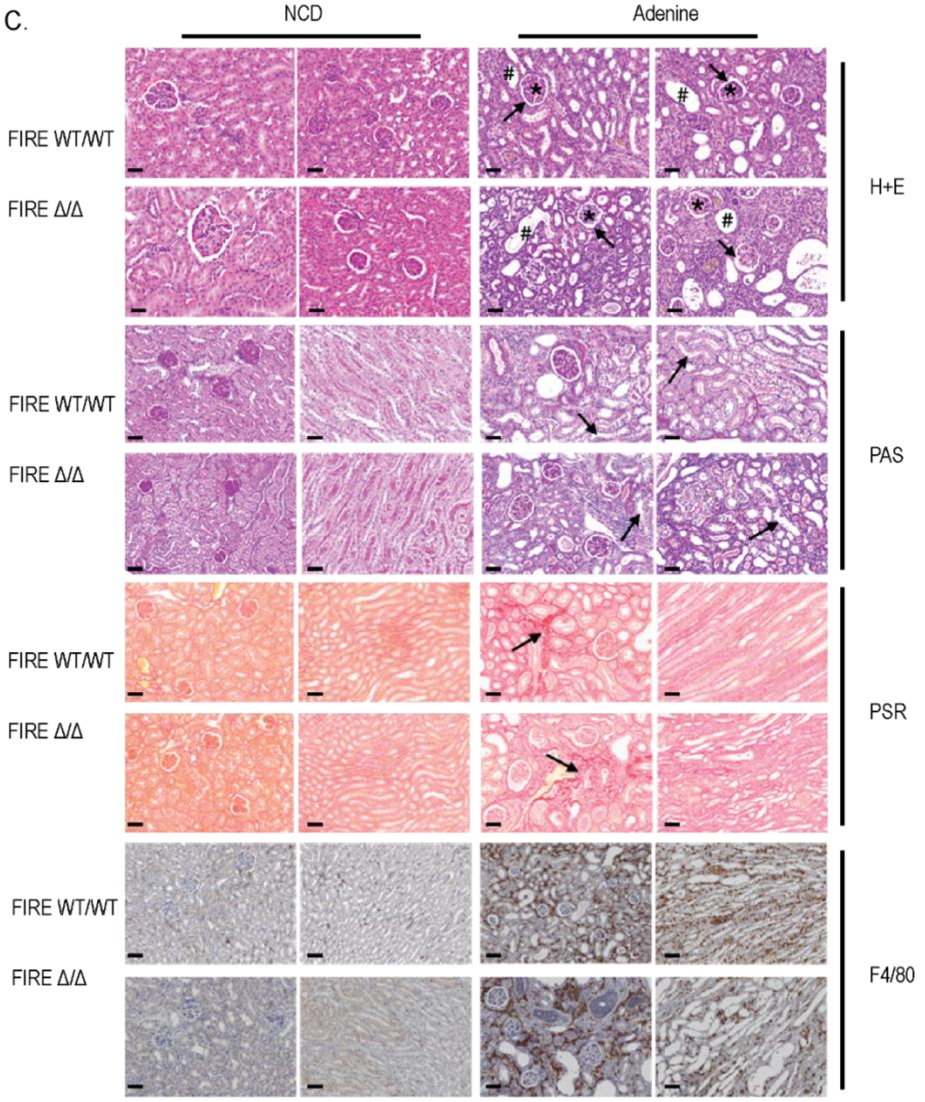


D.

| Parameters | Males | | | | Females | | | |
|---|---|---|---|---|---|---|---|---|
| | Chow | | Adenine | | Chow | | Adenine | |
| | WT | ΔFIRE/ΔFIRE | WT | ΔFIRE/ΔFIRE | WT | ΔFIRE/ΔFIRE | WT | ΔFIRE/ΔFIRE |
| % Total renal affected area | 0.0 | 0.0 | 81.7 ± 6.1 | 81.2 ± 12.5 | 0.0 | 0.0 | 79.4 ± 15.7 | 73.8 ± 6.3 |
| % Tubular atrophy | 0.0 | 0.0 | 35.8 ± 10.2 | 40.6 ± 5.0 | 0.0 | 0.0 | 55.6 ± 21.6 | 57.5 ± 9.6 |
| % Tubules with Casts and cell debris | 0.0 | 0.0 | 23.3 ± 12.1 | 25.0 ± 13.9 | 0.0 | 0.0 | 8.9 ± 5.6 | 7.5 ± 2.9 |
| Apoptosis tubular epithelium | 0.0 | 0.0 | 1.3 ± 0.5 | 1.2 ± 0.5 | 0.0 | 0.0 | 1.6 ± 0.5 | 1.2 ± 0.5 |
| Glomerular sclerosis | 0.0 | 0.0 | 0.0 ± 0.0 | 0.0 ± 0.0 | 0.0 | 0.0 | 0.0 ± 0.0 | 0.2 ± 0.5 |
| Thickening bowman's capsule | 0.0 | 0.0 | 21.7 ± 13.3 | 18.8 ± 11.3 | 0.0 | 0.0 | 0.0 ± 0.0 | 0.2 ± 0.5 |

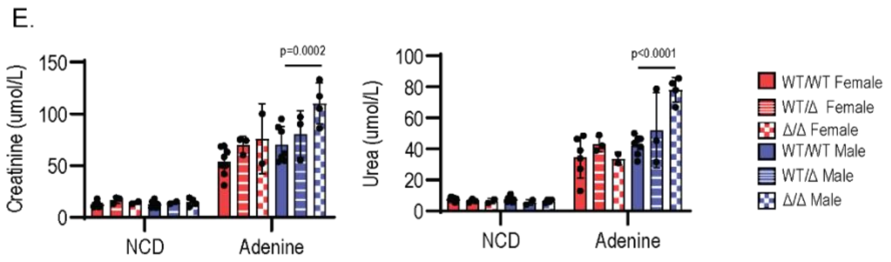

**Figure S8. Adenine-induced chronic kidney disease in *Fireko* mice.** Male and female WT, *Csf1r*$^{\Delta FIRE/+}$ and *Fireko* (FIRE $\Delta/\Delta$) mice were fed normal chow (NCD) or a diet containing 0.2% adenine for 3 weeks. (A) Body weight loss during the treatment period. (B) Kidney weight at sacrifice. (C) Representative sections from male mice (N=5/group) from mice of indicated genotype and diet stained for H&E, periodic acid Schiff (PAS) stain, picrosirius red (PSR) or immunostained for F4/80. (D) Kidney pathology was scored from the H&E sections by a veterinary pathologist (RA) who was blinded to the subject identity as described in Materials and Methods. (E) Serum creatinine and urea. Each point is an individual animal. Data show mean and standard deviation. P values based on 2 way ANOVA with Tukey's multiple comparisons test.

Figure S9

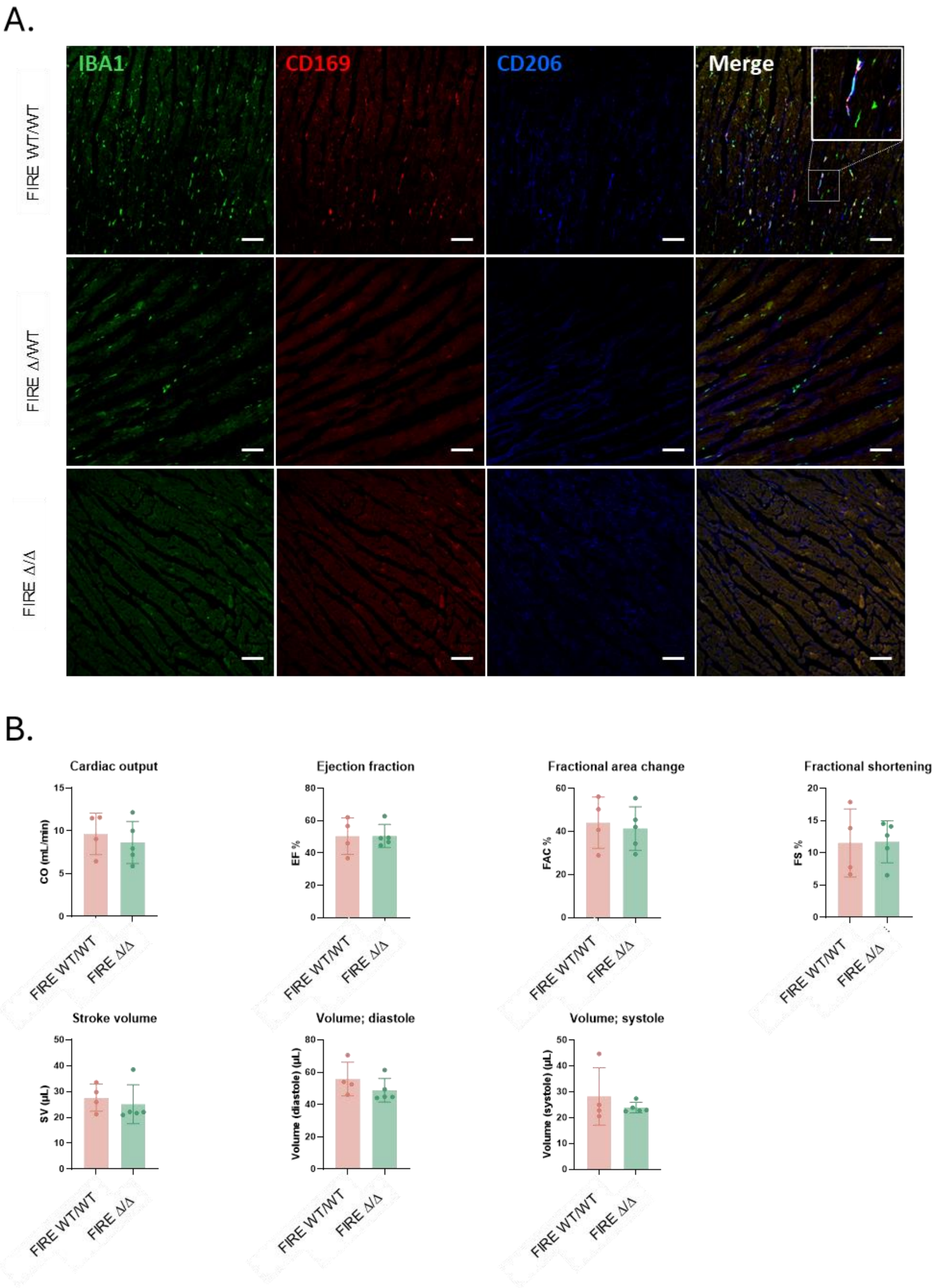


**Figure S9. Cardiac macrophage deficiency in *Fireko* mice does not impact steady state heart function**

(A) Representative images of heart sections from WT, *Csf1r*$^{\Delta FIRE/+}$ and *Fireko* mice at 30 wks of age stained for IBA1, CD169 and CD206 as indicated. (B) Cardiac function was assisted in a cohort of adult male (13-19 wks) WT and *Fireko* mice. Each point is an individual animal.

### Figure S10

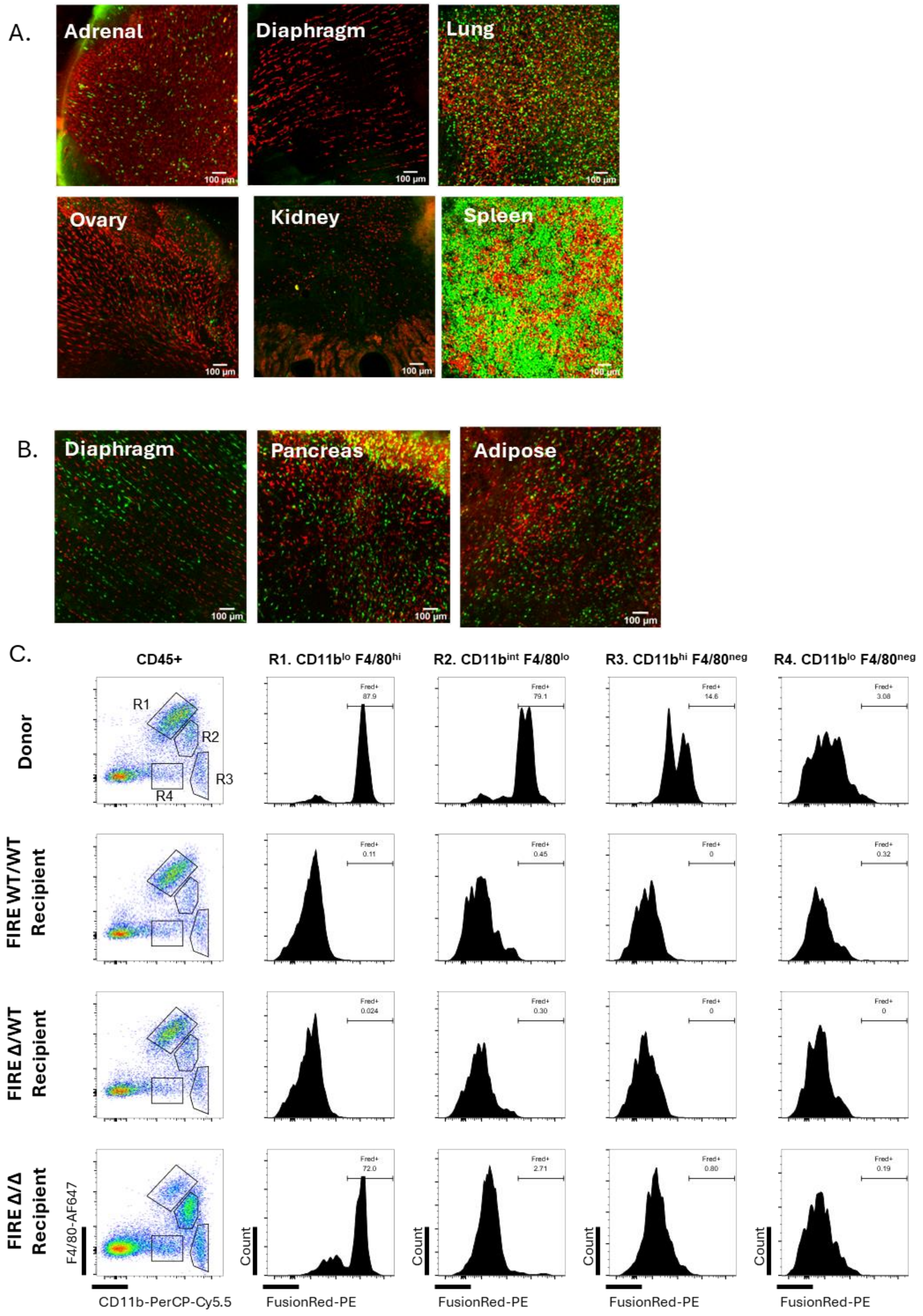
A.
Adrenal
Diaphragm
Lung
Ovary
Kidney
Spleen
100 µm
B.
Diaphragm
Pancreas
Adipose
C.
CD45+
R1. CD11b$^{lo}$ F4/80$^{hi}$
R2. CD11b$^{int}$ F4/80$^{lo}$
R3. CD11b$^{hi}$ F4/80$^{neg}$
R4. CD11b$^{lo}$ F4/80$^{neg}$
Donor
FIRE WT/WT Recipient
FIRE Δ/WT Recipient
FIRE Δ/Δ Recipient
R1
R2
R3
R4
Fred+
87.9
79.1
14.6
3.08
0.11
0.45
0
0.32
0.024
0.30
0
0
72.0
2.71
0.80
0.19
F4/80-AF647
CD11b-PerCP-Cy5.5
Count
FusionRed-PE

**Figure S10. Repopulation of tissue macrophages in *Fireko* mice.**

*Csf1r*-EGFP$^+$ WT, *Csf1r*$^{\Delta FIRE/+}$ and *Fireko* mice were injected i.p. at weaning with WT *Csf1r*-FRed donor cells and chimerism was assessed by whole mount imaging 12 wks post transfer. (A) Representative whole mount images of the tissues indicated in *Fireko* recipients.

(B) Whole mount images of tissues of *Csf1r*$^{\Delta FIRE/+}$ recipients in which donor chimerism was detected.

(C) Representative flow cytometry profiles of kidney macrophage populations in *Csf1r*-FRed donors and WT, *Csf1r*$^{\Delta FIRE/+}$ and *Fireko* bone marrow recipients.
